# Modification of social cheating facilitates the stabilization and chronic infection of polymorphic *Pseudomonas aeruginosa* population

**DOI:** 10.1101/2023.03.08.531804

**Authors:** Kelei Zhao, Xiting Yang, Qianglin Zeng, Yige Zhang, Heyue Li, Jing Shirley Li, Huan Liu, Liangming Du, Yi Wu, Gui Huang, Ting Huang, Yamei Zhang, Hui Zhou, Xinrong Wang, Yiwen Chu, Xikun Zhou

## Abstract

Chronic infection of the common bacterial pathogen *Pseudomonas aeruginosa* frequently leads to the coexistence of heterogeneous individuals to engage in several group behaviors. However, further evolution of the polymorphic *P. aeruginosa* population, including the dynamic change of social cooperation and its impact on host immune system, still remain elusive. We show that the evolution of *P. aeruginosa* in the patients with chronic obstructive pulmonary disease frequently selects the isolates deficient in producing the costly and sharable extracellular products for nutrient acquisition. The evolution of polymorphic *P. aeruginosa* population is mainly concentrated on modifying the adaptability of *lasR* mutants, which are typical cheaters in the competition of quorum-sensing-controlled extracellular proteases. Importantly, *lasR* mutants with varying degrees of evolution interact with wild-type *P. aeruginosa* in a framework termed cascaded public goods game to compete for extracellular proteases and siderophores, and thus perpetuate social cooperation under different conditions. Finally, we find that a polymorphic population comprised of *lasR-*intact *P. aeruginosa* and evolved *lasR-*mutant can minimize the host immune fluctuation for persistent colonization. This study demonstrates the multistage evolution and complex interaction of *P. aeruginosa* in adaptation to host lungs, and provides an explanation for the success of cooperation in public goods game.

## Introduction

Microbes living on the planet evolve diverse capacities to colonize virtually all ecosystems and reserve the possibilities to transfer into new habitats, including human hosts. As a kind of unicellular organism with simple cell structure, the success of bacterial colonization in different environments is largely attributed to the execution of group behaviors, typically social cooperation (1, 2). Bacterial cooperative interactions are mainly mediated by costly and sharable extracellular products (public goods), and can unite the local individuals to increase population fitness in specific habitats (3, 4).

**Figure 1.**
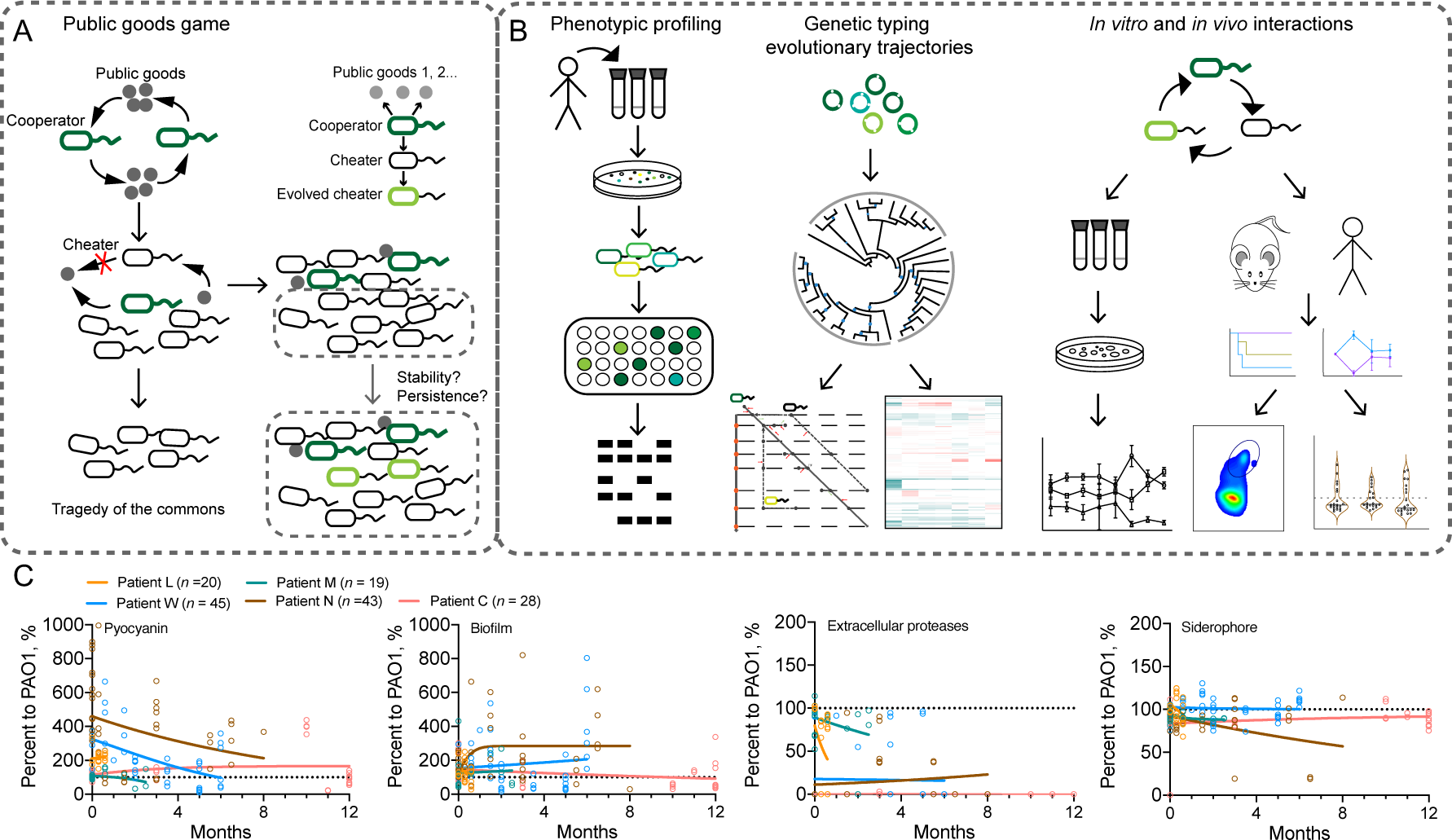
Experimental design and phenotypic changes of *P. aeruginosa* isolates longitudinally collected from host lungs. (*A*) Theoretical model of bacterial public goods game. Public good-producers (cooperators) can be invaded by the nonproducers (cheaters), and the enrichment of nonproducers may cause the tragedy of the commons. We further hypothesize that the evolution of cheaters may lead to a transition of public goods game and facilitate the stabilization and persistent colonization of polymorphic population. (*B*) Experimental design. *P. aeruginosa* isolates were longitudinally collected from the respiratory samples of COPD patients, followed by characterizing their phenotypic and genetic features, phylogenetic status, evolutionary trajectories, and the outcomes of social interactions *in vitro* and *in vivo*. (*C*) Sampling period-dependent changes of the capacities of *P. aeruginosa* COPD isolates to produce extracellular products. Data shown are the phenotypic values of each isolate (symbols) normalized to those of the reference strain PAO1 and the variation tendency (trendlines) in each patient.

**Table 1.** Summary of mutation numbers in evolved P. aeruginosa COPD isolates compared to corresponding initial isolates.

| Sample | Total InDel | CDS InDel | Intergenic InDel | High effect InDel | Total SNP | Syn SNP | Nonsyn SNP | Total SNP | CDS Intergenic SNP | High effect SNP |
| --- | --- | --- | --- | --- | --- | --- | --- | --- | --- | --- |
| W1b | 8 | 3 | 5 | 0 | 23 | 2 | 9 | 11 | 12 | 0 |
| W1c | 360 | 51 | 309 | 20 | 54,644 | 37,152 | 9,708 | 46,902 | 7,742 | 53 |
| W2a | 10 | 4 | 6 | 0 | 12 | 1 | 10 | 12 | 0 | 1 |
| W2b | 9 | 3 | 6 | 0 | 7 | 1 | 5 | 6 | 1 | 0 |
| W2c | 8 | 4 | 4 | 1 | 17 | 3 | 10 | 15 | 2 | 2 |
| W3a | 362 | 50 | 312 | 14 | 54,045 | 36,890 | 9,613 | 46,538 | 7,507 | 45 |
| W4a | 361 | 50 | 311 | 20 | 54,644 | 37,152 | 9,707 | 46,901 | 7,743 | 53 |
| W4b | 10 | 4 | 6 | 0 | 17 | 2 | 5 | 7 | 10 | 0 |
| W4c | 11 | 4 | 7 | 1 | 9 | 0 | 5 | 5 | 4 | 0 |
| W5a | 8 | 3 | 5 | 0 | 9 | 2 | 5 | 7 | 2 | 0 |
| W5b | 366 | 51 | 315 | 15 | 54,030 | 36,889 | 9,612 | 46,537 | 7,493 | 46 |
| W5c | 7 | 3 | 4 | 0 | 23 | 4 | 9 | 14 | 9 | 1 |
| W6a | 7 | 3 | 4 | 0 | 30 | 6 | 7 | 14 | 16 | 1 |
| W6b | 364 | 52 | 312 | 21 | 54,639 | 37,144 | 9,711 | 46,897 | 7,742 | 53 |
| W6c | 10 | 4 | 6 | 0 | 7 | 1 | 5 | 6 | 1 | 0 |
| W7d | 388 | 61 | 327 | 20 | 55,890 | 37,850 | 9,964 | 47,863 | 8,027 | 59 |
| W7e | 9 | 3 | 6 | 0 | 10 | 2 | 6 | 9 | 1 | 1 |
| W7f | 364 | 51 | 313 | 15 | 54,031 | 36,889 | 9,612 | 46,537 | 7,494 | 46 |
| W8a | 10 | 3 | 7 | 0 | 16 | 4 | 8 | 13 | 3 | 1 |
| M2a | 1 | 0 | 1 | 0 | 3 | 2 | 1 | 3 | 0 | 0 |
| M3a | 1 | 0 | 1 | 0 | 18 | 8 | 6 | 14 | 4 | 0 |
| M4a | 1 | 0 | 1 | 0 | 12 | 5 | 1 | 7 | 5 | 1 |
| M5a | 2 | 1 | 1 | 1 | 37 | 2 | 2 | 4 | 33 | 0 |
| L2a | 34 | 5 | 29 | 1 | 29,004 | 18,616 | 5,856 | 24,520 | 4,484 | 48 |
| L2b | 34 | 5 | 29 | 1 | 28,997 | 18,609 | 5,858 | 24,515 | 4,482 | 48 |
| L3a | 34 | 5 | 29 | 1 | 29,008 | 18,611 | 5,864 | 24,523 | 4,485 | 48 |
| L3b | 34 | 5 | 29 | 1 | 28,997 | 18,606 | 5,861 | 24,515 | 4,482 | 48 |
| C5a | 31 | 5 | 26 | 3 | 33,170 | 21,769 | 6,716 | 28,547 | 4,625 | 44 |

According to the theory of the classic public goods game, cooperation is vulnerable to the sabotage of cheaters who do not produce public goods but benefit from the cooperation of producers (cooperators). Cheating as the optimal strategy will ultimately overtake cooperation and results in the tragedy of the commons due to the shortage of crucial public goods (5, 6) (Fig. 1*A*). However, population collapse rarely occurs during the colonization of bacterial populations in natural environments or in host tissues and is only observed when the growth of bacterial cells is determined by a single kind of public good under laboratory conditions. All these cases can be supported by the studies focusing on the phenotypic and genetic evolution of *Pseudomonas aeruginosa*, which is a ubiquitous Gram-negative opportunistic bacterium capable of colonizing a wide range of natural and clinical environments (3, 7–10).

The large genome size and complicated intracellular transcriptional network of *P. aeruginosa* endow the bacterium with the capacity to engage in a variety of cooperative interactions, such as the extracellular elastase-mediated acquisition of macromolecular nutrients, extracellular siderophore-mediated chelation of irons and exopolysaccharide-related biofilm formation (7, 11–13). Laboratorial evidence has confirmed the massive invasion of wild-type (WT) *P. aeruginosa* by the strains deficient in producing a single extracellular product, which is vitally required in each scenario of the public goods game. By contrast, these mutants are rare in natural environments but abundant in the lungs of immunocompromised patients (8, 14, 15). Colonization of *P. aeruginosa* in host lungs may encounter various environmental pressures, including low nutrient availability, inflammatory responses, antibiotic clearance, and inter/intraspecific competitions. These pressures may pose a metabolic burden on *P. aeruginosa* to produce the corresponding public goods and thereby lead to the enrichment of cheaters with a specific mutation during further evolution (16–19).

Among *P. aeruginosa* mutants from the lungs of patients with the genetic disease cystic fibrosis (CF), loss-of-function mutations in the *lasR* gene, which encodes the central regulator of the quorum-sensing (QS) system and positively controls the production of a series of extracellular proteases, are the most frequent (14, 20). The *P. aeruginosa* QS system comprises three core regulators, and only the *lasR* mutant can invade QS-intact individuals in a cheating manner under the condition that the QS-controlled public goods are the key limiting factor for individual growth (21). Additionally, CF airways are frequently colonized by a mixture of *P. aeruginosa* isolates with intact and mutated *lasR* genes, rather than by a predominating frequency of *lasR* mutants (cheaters) according to the theory of public goods game (14, 22, 23). Several empirical and mathematical studies have provided conceptual explanations for the maintenance of cooperation in the *P. aeruginosa* population, such as social policing, metabolic prudence, and cheating on cheaters (17, 24–26). Nevertheless, the cooperative interactions of *P. aeruginosa in vivo* might be multifactorial and more complicated. Few studies have explored the evolutionary dynamics of *P. aeruginosa* population comprises *lasR-*intact and *lasR* mutant individuals in host lungs. As *lasR* mutant cheaters have a competitive advantage over WT cooperators and some evolved *lasR* mutants can also elicit hyperinflammatory responses (7, 27), we hypothesize that further evolutionary selection in the polymorphic *P. aeruginosa* population may mainly focus on modifying the adaptability of *lasR* mutants to facilitate the persistent colonization of the whole population (Fig. 1*A*).

In this study, by longitudinally collecting *P. aeruginosa* isolates from the bronchoalveolar lavage (BAL) fluids of patients with chronic obstructive pulmonary disease (COPD), followed by phenotypic, genetic, phylogenetic, and multi-omics-based functional analyses (Fig. 1*B*), we found that the evolution of *P. aeruginosa* in COPD airways frequently selects isolates deficient in producing the costly and sharable extracellular products for nutrient acquisition. The *in vivo* evolution of polymorphic *P. aeruginosa* was characterized by modifying the adaptability of *lasR* mutants, as determined by the evolutionary trajectory analysis. We then studied the relationship of *lasR* mutants with varying degrees of evolution and the ancestral *P. aeruginosa* under different conditions and identified an interaction of the cascaded public goods game. Finally, the contribution of evolved *lasR-*mutant to the survival advantage of *P. aeruginosa* population was verified by using mouse models and clinical respiratory samples.

## Results

### *In vivo* evolution of *P. aeruginosa* selects individuals deficient in producing sharable products for nutrient acquisition

To explore the social traits of *P. aeruginosa* in host lungs, 25 *P. aeruginosa*-positive COPD patients (but negative in the past whole year) were enrolled for the longitudinal collection of *P. aeruginosa*. BALs or spontaneous sputa of the patients were recovered at intervals of approximately 15 days, and a total of 536 *P. aeruginosa* colonies were obtained. The patients could be divided into three groups according to their lifetime after the first sampling: Group A included fourteen patients who passed away within 0.5 to 2.5 months because of severe lung infection, Group B included three patients who lived for 4 to 8 months, and Group C included only one patient who became *P. aeruginosa* negative after 12 months. The remaining seven patients quit the project after 1 or 2 sampling periods because of remission and other uncontrollable factors. *P. aeruginosa* isolates of patient L (80 years old, *n* = 20 in 0.6 months) and patient M (74 years old, *n* = 19 in 2.5 months) from Group A, patient W (92 years old, *n* = 45 in 6 months) and patient N (81 years old, *n* = 43 in 8 months) from Group B, and patient C (66 years old, *n* = 28 in 12 months) from Group C with complete sampling periods were selected for further analyses.

Among the virulence-related phenotypes, we mainly evaluated the levels of extracellular products (including pyocyanin, biofilm, QS-controlled extracellular proteases, and siderophores) produced by *P. aeruginosa* isolates of each patient as the sampling periods increase. The results showed that the majority of isolates from each patient throughout the sampling periods produced higher or comparable levels of pyocyanin and biofilms to the reference strain *P. aeruginosa* PAO1. In contrast, the production of extracellular proteases and siderophores, especially the former by the COPD isolates, was lower than those of PAO1 and gradually reduced in patients L and M, while it remained on the low side by the isolates of the other three patients (Fig. 1*C*). Moreover, all the *P. aeruginosa* isolates of patient C were deficient in producing QS-controlled extracellular proteases, and this patient was still alive after the sampling was finished. These results suggested that QS-controlled extracellular proteases might be a vitally important public goods that could bring more direct benefits (in terms of nutrient acquisition) for the survival of *P. aeruginosa in vivo*, followed by siderophores, and the intraspecific competitions of which would be more readily to cause the invasion of individuals deficient in producing the products.

### COPD airways are frequently colonized by polymorphic *P. aeruginosa* population

The genetic relationships of the clinical *P. aeruginosa* isolates were preliminarily sorted by enterobacterial repetitive intergenic consensus-polymerase chain reaction (ERIC-PCR)-based isolate typing. The results revealed that *P. aeruginosa* isolates longitudinally collected from the same COPD patient could be generally separated into two or three subgroups (Fig. S1), indicating the colonization of multiclonal *P. aeruginosa*. To evaluate the mutation status and relationship of *P. aeruginosa* from COPD airways, the isolates representing the phenotypes of other isolates in each sampling period from the corresponding patients and those distributed in the main branches of isolate typing were selected for whole-genome sequencing (WGS)-based comparative genomic analyses.

The results of WGS showed that the genome size of the sequenced *P. aeruginosa* isolates ranged from 6.19 to 6.74 Mbp (6.49 ± 0.116 Mbp), with an average GC content of 66.38% and gene number of 5977 ± 121.20 (Table S1). We then identified the mutations that arose in *P. aeruginosa* during evolution by comparing the genome sequences of the evolved isolates to those of the corresponding initial isolates. As shown in Table 1, the isolates obtained from patient L in the two sampling periods had almost the same numbers of insertions and deletions (InDels) and single nucleotide polymorphisms (SNPs). The isolates from patient M harbored 1-2 InDels and 3-37 SNPs. Isolate C5a, which evolved for 12 months in the lungs of patient C, harbored 31 InDels and 33,170 SNPs. The isolates from patient W could be separated into two groups according to the number of mutation sites. The isolates deficient in producing QS-controlled extracellular proteases accumulated considerably larger numbers of mutations harmful to gene function than those of extracellular protease-positive isolates (Tables 1, S2, and S3).

SNP-based phylogenetic analysis further revealed that *P. aeruginosa* isolates sequenced in this study could be separated into three main groups by also considering their multilocus sequencing typing (MLST) and serotyping results (Fig. 2*A*). Specifically, Group 1 was composed of 13 *P. aeruginosa* isolates belonging to MLST type ST357 and serotype O11 (ST357:O11) obtained from almost all sampling periods of patient W. Group 2 was composed of all the isolates from patient M, three isolates from patient W, and isolate C1a from patient C. These isolates belonged to the same MLST and serotype as ST233:O6. The isolates in Group 2 showed a close relationship with the hypervirulent clinical isolate *P. aeruginosa* PA14. The composition of the *P. aeruginosa* isolates in Group 3 was complex in terms of their isolation source, MLST, and serotype, including all the isolates from patient L (ST549:O5 and ST181:O3), four isolates from patient W (ST1129:O6 and ST549:O3), and isolate C5a from patient C (ST274:O3). These isolates were clustered with the model *P. aeruginosa* isolates PAK, PAO1, and CF isolate DK2. Notably, the lungs of patient W, who was treated in the intensive care unit throughout the sampling period, were colonized by *P. aeruginosa* isolates with four kinds of MLST:serotype combinations. These isolates were distributed in all three phylogenetic groups and clustered with isolates from other patients. Moreover, patients L and C were also infected by *P. aeruginosa* isolates with two kinds of MLST:serotype combinations. Therefore, our results clearly demonstrated the cross-transmission of *P. aeruginosa* among patients and confirmed that COPD airways were frequently cocolonized by polymorphic *P. aeruginosa*.

**Figure 2.**
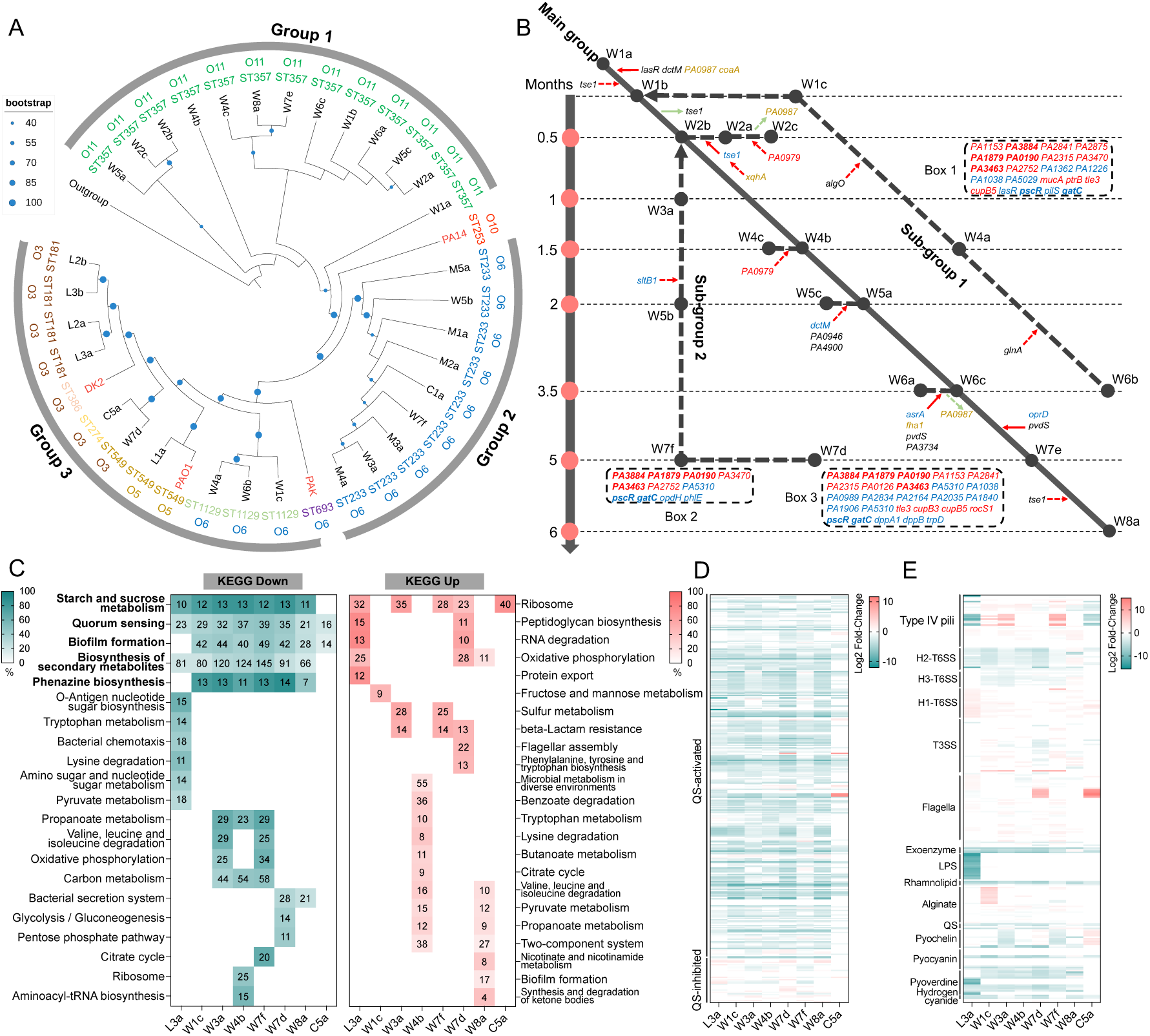
Phylogenetic analysis and evolutionary trends of *P. aeruginosa* isolates in host lungs. (*A*) SNP-based phylogenetic tree of *P. aeruginosa* COPD isolates constructed using maximum likelihood method. The size of blue dot at each node indicates bootstrap value (1000 replicates). All the isolates can be generally separated into three groups. The colored outer ring indicates MLST and serotypes. Outgroup, *Azotobacter vinelandii*. (*B*) Evolutionary trajectories of *P. aeruginosa* isolates in patient W. Time-dependent accumulation of mutation sites in the genes of *P. aeruginosa* isolates from patient W were determined by comparing the whole-genome sequence of each isolate to W1a. Red solid arrow, acquired fixed mutation. Red dotted arrow, acquired accidental mutation. Green solid arrow, lost fixed mutation. Green dotted arrow, lost accidental mutation. Blue color, gene with high effect SNP. Red color, gene with frame-shifted InDel. Black color, gene with nonsynonymous SNP. Yellow color, gene with non-shifted InDel. Bold, mutated genes shared by the subgroups. (*C*) KEGG terms enriched by the significantly down-and up-regulated genes of evolved *P. aeruginosa* COPD isolates compared to the corresponding initial isolates (*P* < 0.05). Heatmap color indicates percentage of differentially express gene number (black number in the colored squares) to the background gene number in each KEGG term. (*D*) Expression changes of QS-controlled genes (315 QS-activated and 38 QS-inhibited) in evolved *P. aeruginosa* COPD isolates compared to the corresponding initial isolates. (*E*) Expression changes of virulence-related genes in evolved *P. aeruginosa* COPD isolates compared to the corresponding initial isolates.

### *In vivo* evolution of polymorphic *P. aeruginosa* is concentrated on modifying the adaptability of *lasR* mutants

To determine the genetic features of *P. aeruginosa* that were associated with the differentiation of social behaviors and the evolution of polymorphic *P. aeruginosa* in the lung environments, the evolutionary trajectories of *P. aeruginosa* isolates from patient W were predicted due to their dominant phylogenetic status and abundant genetic diversity (Fig. 2*A*). As shown in Fig. 2*B*, W1a was identified as the primary colonizing clone because it was initially isolated from the patient and had an intact virulence repertoire compared to PAO1. All the isolates with the same MLST and serotype of ST357:O11 distributed in Group 1 of the phylogenetic tree were defined as the main evolutionary group (Figs. 2*A* and *B*). W1c, W4a, and W6b, with the MLST and serotype of ST1129:O6, clustered together in Group 3 of the phylogenetic tree and were defined as Subgroup 1. Finally, isolates W3a, W5b, and W7f, with the MLST and serotype of ST233:O6, clustered together in Group 2 of the phylogenetic tree were defined as Subgroup 2.

Compared to W1a, all the core isolates in the main group harbored 5 nonsynonymous SNPs in *lasR* and *dctM* (C4-dicarboxylate transporter) and 3 non-shifted InDels in *PA0987* and *coaA* (Figs. 2*B*, *C*, and S2). Specifically, only W1b harbored an additional nonsynonymous SNP in the type VI secretion system (T6SS) effector-encoding gene *tse1*. W2a harbored 1 nonsynonymous and 1 premature stop SNP in *tse1* and 1 non-shifted InDel in *xqhA* (T2SS secretion protein), and the premature stop SNP in *tse1* was passed on to W2c. W2c lost the parental non-shifted InDel in *PA0987* but harbored a new frame-shifted InDel in *PA0979*. W4c acquired an additional shifted InDel in *PA0979* compared to W4b. W5c gained 7 SNPs, including 1 premature stop and 6 nonsynonymous mutations in *dctM*, *PA0946*, and *PA4900*, and lost 1 non-shifted InDel site in *coaA* and 2 nonsynonymous SNPs in *dctM*. W6a lost the non-shifted InDel in *PA0987* and harbored 1 non-shifted InDel in *fha1* (T6SS secretion protein) and 3 new SNPs. Compared to other core isolates, 1 premature stop SNP in *oprD* (multifunctional outer membrane porin) and 1 nonsynonymous SNP in *pvdS* (pyoverdine biosynthesis regulatory gene) were fixed in W7e and W8a, and W8a accumulated another nonsynonymous SNP in *tse1*. In Subgroup 1, W1c, W4a, and W6b carried premature stop SNP sites in *lasR* and showed the same categories of mutated genes, including *pscR*, *pilS*, and *gatC* with premature stop SNP sites, *mucA*, *tle3*, and *cupB5* with frame-shifted InDel sites, and other unknown genes (Fig. 2*B*-Box 1). W4a harbored a new nonsynonymous SNP in *algO*, and W6b harbored a new nonsynonymous SNP in *glnA*. In Subgroup 2, W3a, W5b, and W7f showed the same categories of genes with mutation sites potentially harmful to gene function (Fig. 2*B*-Box 2), and the majority of the common genes were transmitted to W7d (Fig. 2*B*-Box 3). These results indicated that the evolution of polymorphic *P. aeruginosa* population in COPD airways might mainly concentrate on modifying the adaptability of *lasR* mutants, especially the secretion and regulation of extracellular products.

To investigate the effect of accumulated mutations on the functional categories of *P. aeruginosa*, the global transcriptions of *P. aeruginosa* isolates from the initial and final sampling periods and those from the subgroups of patient W were profiled by RNA sequencing (RNA-seq). Compared to the corresponding initial isolates, the results of KEGG pathway prediction demonstrated the greatest decreases in functional categories related to QS, followed by starch and sucrose metabolism, biofilm formation, biosynthesis of secondary metabolites, and phenazine biosynthesis in all the evolved *P. aeruginosa* isolates (Fig. 2*C* and Dataset S1). Interestingly, the 5 commonly decreased KEGG terms were the same as those enriched by the genes activated by the two main QS regulator genes, *lasR* and *rhlR* (Dataset S1). When the significantly differentially expressed genes were mapped to the list of QS-controlled (315 activated and 38 inhibited) genes (20), we found that all the isolates showed decreased expression of a large number of genes activated by QS (Figs. 2*D* and S3). Additionally, the significantly enriched KEGG terms differed among the evolved isolates as the sampling time increased (Fig. 2*C*). We further compared the expression levels of all the virulence-related genes in the evolved *P. aeruginosa* to the corresponding initial isolates and found that, the expression levels of genes related to QS-controlled extracellular products, iron acquisition, H2-and H3-T6SS were decreased, while the genes related to H1-T6SS and flagella were generally increased. On the other hand, the expression levels of genes related to type IV pili, T3SS, and alginate production varied among isolates (Fig. 3*E* and Dataset S2). Therefore, these results combined with the generally decreased production of QS-controlled extracellular proteases by *P. aeruginosa* isolates from COPD airways (Fig. 1*C*), revealed that the transcriptional divergence of QS regulation caused by the mutation in the *lasR* gene commonly occurred in evolving polymorphic *P. aeruginosa* population, while the adaptability of *lasR* mutants would be adjusted by the accumulation of other mutations during further evolution in COPD airways.

**Figure 3.**
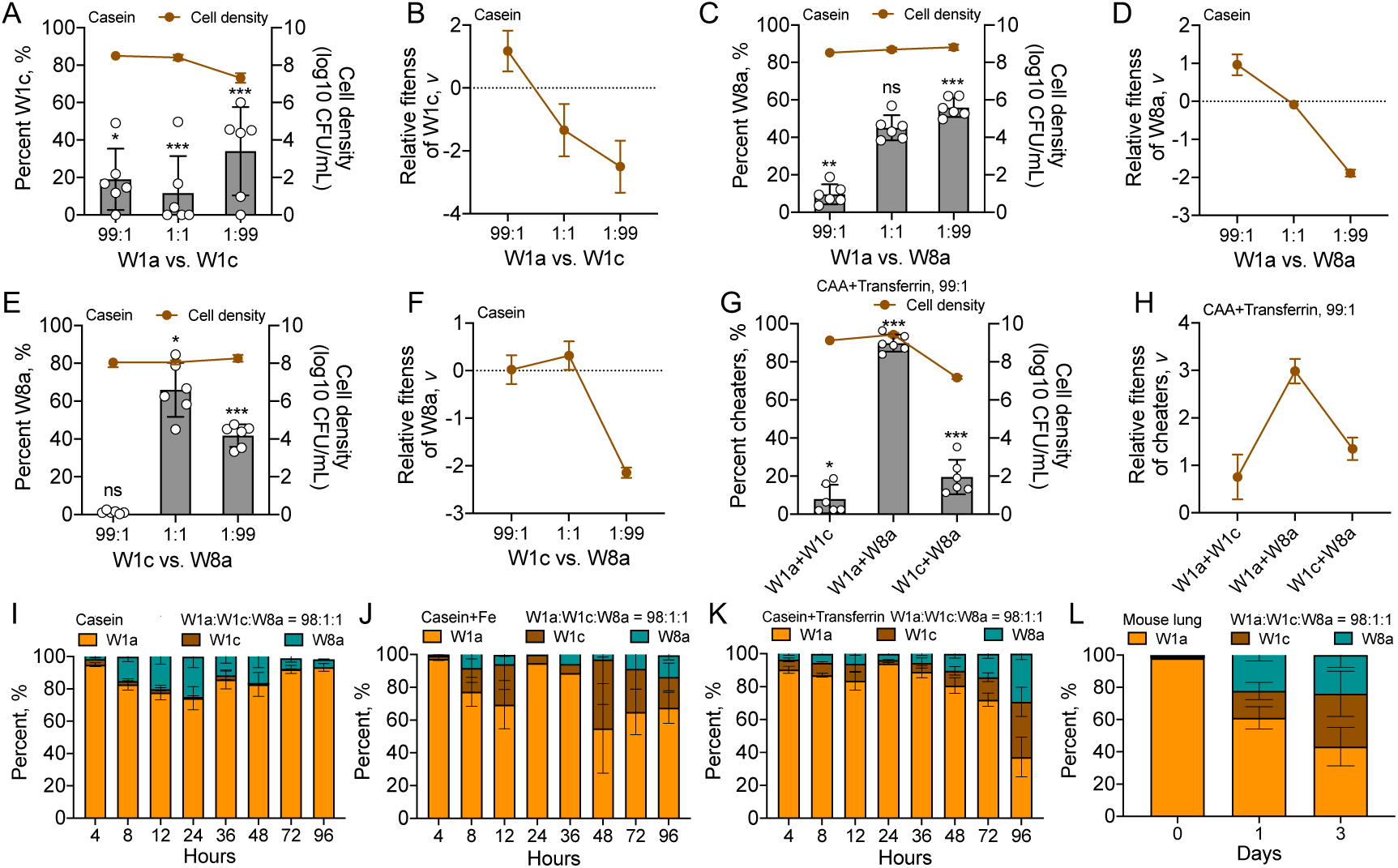
*In vitro* and *in vivo* competitions of successively identified *P. aeruginosa* isolates under different conditions. *P. aeruginosa* isolates W1a (*lasR-*intact), W1c (evolved *lasR* mutant), and W8a (*lasR-pvdS* mutant) were cocultured in double or triple in 2 mL of M9 minimal growth medium supplemented with different carbon sources and iron levels from different initial ratios. Frequencies and relative fitness of W1c or W8a in double cocultures of (*A* and *B*) W1a+W1c, (*C* and *D*) W1a+W8a, or (*E* and *F*) W1c+W8a in M9-casein medium for 24h. (*G*) Frequencies and (*H*) relative fitness of the isolates with an initial frequency of 1% in double cocultures of W1a+W1c, W1a+W8a, or W1c+W8a in iron-limiting M9-CAA medium for 24h. Right Y-axis indicates the cell densities of each culture. The value of each column was compared to the initial frequency of corresponding isolate using two tailed unpaired *t-*test. *, *P* < 0.05. **, *P* < 0.01. ***, *P* < 0.001. Dynamic changes in the frequencies of W1a, W1c, and W8a during the coevolution of the three isolates in (*I*) M9-casein, (*J*) iron-abundant M9-casein, (*K*) iron-limiting M9-casein media, and (*L*) mouse lungs from an initial ratio of 98:1:1. The culture media were refreshed at 24 h interval and the experiment was stopped when the frequency of each isolate in the culture was relatively stable. Data shown are means ± standard deviation (SD) of six independent replicates.

### *lasR* mutant-centered cascaded public goods game stabilizes social cooperation

Based on the genetic and transcriptional characteristics of *P. aeruginosa* isolates identified above (Fig. 2), the isolate W1a from the initial sampling period with intact *lasR* gene, W4b with shifted InDel mutation in *lasR*, W8a with an additional shifted InDel in *pvdS*, and W1c with several loss-of-function mutations in pathoadaptive genes including *lasR*, which were representative of the genetic feature of the others, were selected to study their social interactions in the scenario of public goods game. We first showed that when the isolates were monocultured in the QS-required medium (M9 + casein), the growth of W4b and W8a was slower than that of W1a in the initial 6 h and then became comparable to W1a, while the growth of W1c was remarkable slower than the other isolates (Fig. S4*A*). Correspondingly, W1a produced a large proteolytic ring on M9-casein plates, W4b and W8a produced smaller proteolytic rings, while the capacity of W1c to produce the QS-controlled extracellular proteases was completely abolished (Fig. S4*B*). By contrast, W4b and W8a grew faster than W1a in M9 casamino acids (CAA, hydrolysates of casein) in the initial 12 hours but became slower after 20 hours, and W1c also grew slower than the other isolates (Fig. S4*C*).

We then performed a batch of competition assays to investigate the interactions of the selected *P. aeruginosa* isolates by coculturing different combinations of them in M9-casein medium. The results showed that both the growth of W1c and W8a from a small initial frequency (1%) was faster than that of cocultured W1a in the two-player game (Fig. 3*A-D*), indicative of the exploitation of W1a by W1c and W8a in a cheating manner. W4b as a transitional isolate from W1a to W8a could also invade W1a (Fig. S5). However, the three *lasR* mutants showed different degrees of exploitation on W1a, and the ability of W1a to invade the three *lasR* mutants was also different. The population collapse caused by the enrichment of cheaters was detected only in W1a-W1c coculture group. These differences might be associated with the different capacities of the *lasR* mutants to produce elastase (public goods) and the innate slow growth rate of W1c (Fig. S4). We also tested the interaction of W1c and W8a and found that, W8a failed to invade W1c while W8a could be readily exploited by W1c (Fig. 3*E*, *F*). This was due to the weak ability of W8a to produce costly public goods, while W1c produced nothing (Fig. S4*B*).

Moreover, we found that the isolate W8a, which harbored mutated *pvdS* gene, failed to invade its transitional parental isolate W4b (*pvdS*-intact) from a small initial frequency in M9-CAA medium, but could significantly exploit W4b under iron-depleted conditions that caused by the supplementation of Transferrin (Fig. S6). This result indicated that as two kinds of *lasR* mutants, the latterly identified isolate W8a evolved an additional capacity to cheat on W4b in the public goods game mediated by iron-chelating siderophores. W8a had a remarkably higher relative fitness than W1a in the public goods game of siderophores and could also exploit W1c (Fig. 3*G*, *H*). We then sequenced the *lasR* and *pvdS* genes of all the *P. aeruginosa* isolates from the five COPD patients and found that the emergence of *lasR* mutants was earlier than that of *pvdS* mutants, and about one sixth of the *lasR* mutants from patient W co-carried a mutation in the *pvdS* gene (Fig. S7). Therefore, our results demonstrated the cheating behaviors of different *P. aeruginosa lasR* mutants from the same patients in playing public goods games in dependence on environmental change. Additionally, the extracellular protease-mediated public goods game is more likely to be the main scenario that frequently happens in the *P. aeruginosa* population in the host lung, while the competition of siderophores might only occur during further evolution (Figs. 2*B* and S7).

We further monitored the interaction dynamics of W1a, W1c, and W8a in three-player public goods game by successively coculturing them (98:1:1) in M9-casein medium with different levels of iron. The results showed that in casein medium which creates the scenario of an extracellular protease-mediated public goods game, the frequency of W8a increased to 25.03 (± 6.82) % in the initial 24 h and then gradually decreased to 6.54 (± 2.35) % after 24 h, while W1c showed a weak ability to invade W1a and was gradually eliminated from the game after 48 h (Fig. 3*I*). Interestingly, the addition of iron in M9-casein medium resulted in the success of W1c. This might be partially related to the restored growth of W1c by supplementation with iron (Fig. S8). Both W8a and W1c could invade W1a and the three isolates coexisted with a ratio of approximately 3:1:1 (Fig. 3*J*). In casein+Transferrin medium, which creates the scenario of multiple public goods games, the three isolates could still coexist in the population with a W1a/W1c/W8a ratio of approximately 1:1:1 (Fig. 3*K*). Additionally, none of the three sets of successive subculturing assays incurred a collapse of the population, although the growth of the mixed population was relatively slow in casein+Transferrin medium (Fig. S9*A*). We then tested the interactions of the three isolates in mouse lungs by coinfecting W1a, W1c, and W8a from an initial ration of 98:1:1. The results showed that W1c and W8a could invade W1a and coexist in lung environments (Fig. 3*L* and S9*B*).

These results collectively confirmed that *lasR* mutants with varying degrees of evolutions can interact with WT *P. aeruginosa* in a framework termed cascaded public goods game, which contributes to the maintenance of cooperation and the relative stability of population structure in diverse environments.

### A mixture of WT *P. aeruginosa* and evolved *lasR* mutant restrains host inflammatory responses

Based on the finding above that the evolution of polymorphic *P. aeruginosa* population mainly selected *lasR* mutants that could enhance the survival fitness of the population (Figs. 1*C* and 2), we then set out to test the influence of the invasion of evolved *lasR* mutant on the adaptability (in terms of bacterial virulence and host immune fluctuation) of *P. aeruginosa* population by establishing chronic infection models using W1a, W1c, and a mixture thereof (1:1).

In the slow-killing assay (mimicking the chronic infection status of *P. aeruginosa*) conducted in the *Caenorhabditis elegans* infection model, W1a caused earlier death of the nematodes, but the results did not significantly differ from the mortality of nematodes infected by the mixture of W1a and W1c (*P* = 0.1612). In contrast, the evolved *lasR* mutant W1c failed to kill any nematodes over 144 hours (Fig. S10). Unsurprisingly, compared to the 100% survival rate of mice chronically challenged by W1c, W1a killed 80% of the mice, while W1a-W1c coinfection killed 50% in the initial 4 days (Fig. 4*A*). Moreover, although the growth of W1c was remarkably slower than that of W1a *in vitro*, it could persist in the lungs of mice infected by the mixture of W1a and W1c and showed less fluctuated residual cell numbers in W1c-infected mice (Figs. 4*B*, S4 and S11). These results suggested that W1a had a strong pathogenic ability, while W1c lost most of the lethality but adapted to persistent colonization. Therefore, we suspected that a mixture of W1a and W1c might combine the advantages of the two individuals and facilitate the colonization of *P. aeruginosa* during chronic lung infection.

**Figure 4.**
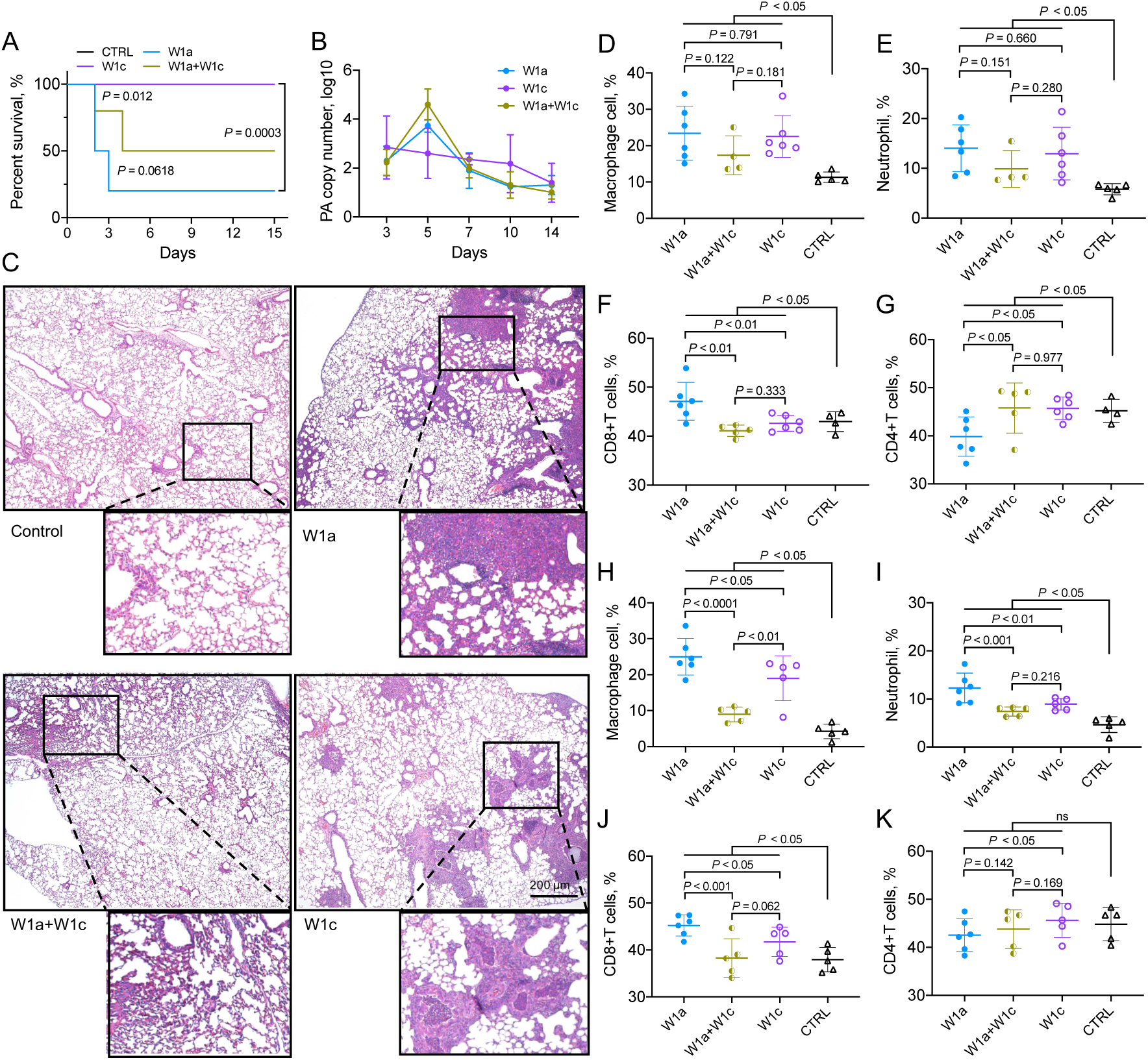
Pathogenicity of *lasR-*intact *P. aeruginosa* and evolved *lasR* mutant in mouse model. (*A*) Chronic lung infection mouse model (40 mice per group) using *P. aeruginosa* COPD isolates W1a, W1c, and a 1:1 mixture of them. The survival curves of mice were compared by using Log-rank (Mantel-Cox) test. (*B*) 16S rRNA-based copy number change of *P. aeruginosa* during chronic infection of mouse lungs. (C-K) Lung tissues of mice chronically infected with/without *P. aeruginosa* at (*C-G*) day 3 and (*H-K*) day 7 were collected. (*C*) Lungs embedded in formalin were evaluated by H&E staining. Images are representative of three independent replicates. Scale bar, 200 μm. (*D-K)* The proportions of inflammatory macrophages, neutrophils, CD8^+^ T cells, and CD4^+^ T cells were evaluated by flow cytometry. 4-6 mice per group. The data shown are the means ± SD of three independent replicates. Statistical significance was calculated using one-way ANOVA with Tukey post-hoc tests.

The results of histological staining revealed that the W1a-W1c coinfection group caused the least inflammatory cell infiltration to the mouse lung at 3 days post infection (Fig. 4*C*), while the W1a and W1c groups showed severe lung damage. We further examined the change of immune status in lung tissues induced by different combinations of *P. aeruginosa* during chronic infection by flow cytometry. Accordingly, the W1a group recruited the most inflammatory macrophages, neutrophils, and CD8^+^ T cells, while the W1a-W1c coinfection group recruited the least (Figs. 4*D-F* and S12). Notably, the lung tissues had the lowest level of CD4^+^ T cells in the W1a group, and there were no significant changes in CD4^+^ T cells in the W1a-W1c coinfection and W1c groups compared to the counterparts of the control group. This indicated that the mice of W1a-infected group had the most significant immune fluctuations on day 3 (Figs. 4*G* and S12*C*). Furthermore, the levels of immune cells in the lungs of these three groups on day 7 were similar to those comparts on day 3, except for CD4^+^ T cells, which were no longer significantly different among the different groups (Figs. 4*H-K* and S13). These above results indicated that the interactions between W1a and W1c had a significant impact on host defense immunity. The evolved *lasR* mutant might help the WT isolate to impair host immune activation and thus facilitates chronic *P. aeruginosa* infection.

### Evolved polymorphic *P. aeruginosa* population compromises host inflammatory gene expression

To better understand the effects of the polymorphic *P. aeruginosa* population on host immune responses, RNA-seq was then performed using mouse lung tissues above (Fig. 4*B*). At day 3 post infection, all the three infection groups triggered significant transcriptional changes in mouse lungs compared to the uninfected group. The most enriched upregulated signaling pathways were cytokine-cytokine receptor interaction, IL-17, TNF, NF-κB, and Nod-like receptor signaling pathways, etc. (Fig. 5*A*). However, the fold-change of differently expressed genes was significantly different among the three groups. The W1c group had the most significantly changed genes related to the host immune response, while the W1a-W1c coinfection group had the fewest (Fig. 5*A*). We further divided these significantly changed genes into three categories: immunomodulatory molecules, cytokine-receptor interaction, and ECM (extracellular matrix)-receptor interaction. The expression of many representative molecules involved in host immune defense processes, such as Mevf, IL23a and Mmp9, had the smallest fold-change in the W1a-W1c coinfection group (Figs. 5*B-D* and S14*A-C*). Consistently, the W1a-W1c coinfection group also induced the slightest dynamic transcriptional change in mice compared with the other two groups between day 3 and day 7 (Figs. 5*E* and S14*D*). These results indicated that *P. aeruginosa* W1a alone had the strongest pathogenicity, whereas W1c might have lost most of its virulence and passively survived in the host by regulating its own transcriptional state. Therefore, the polymorphic *P. aeruginosa* population comprised of QS-intact and -deficient individuals had a unique advantage in persistently colonizing host lungs.

**Figure 5.**
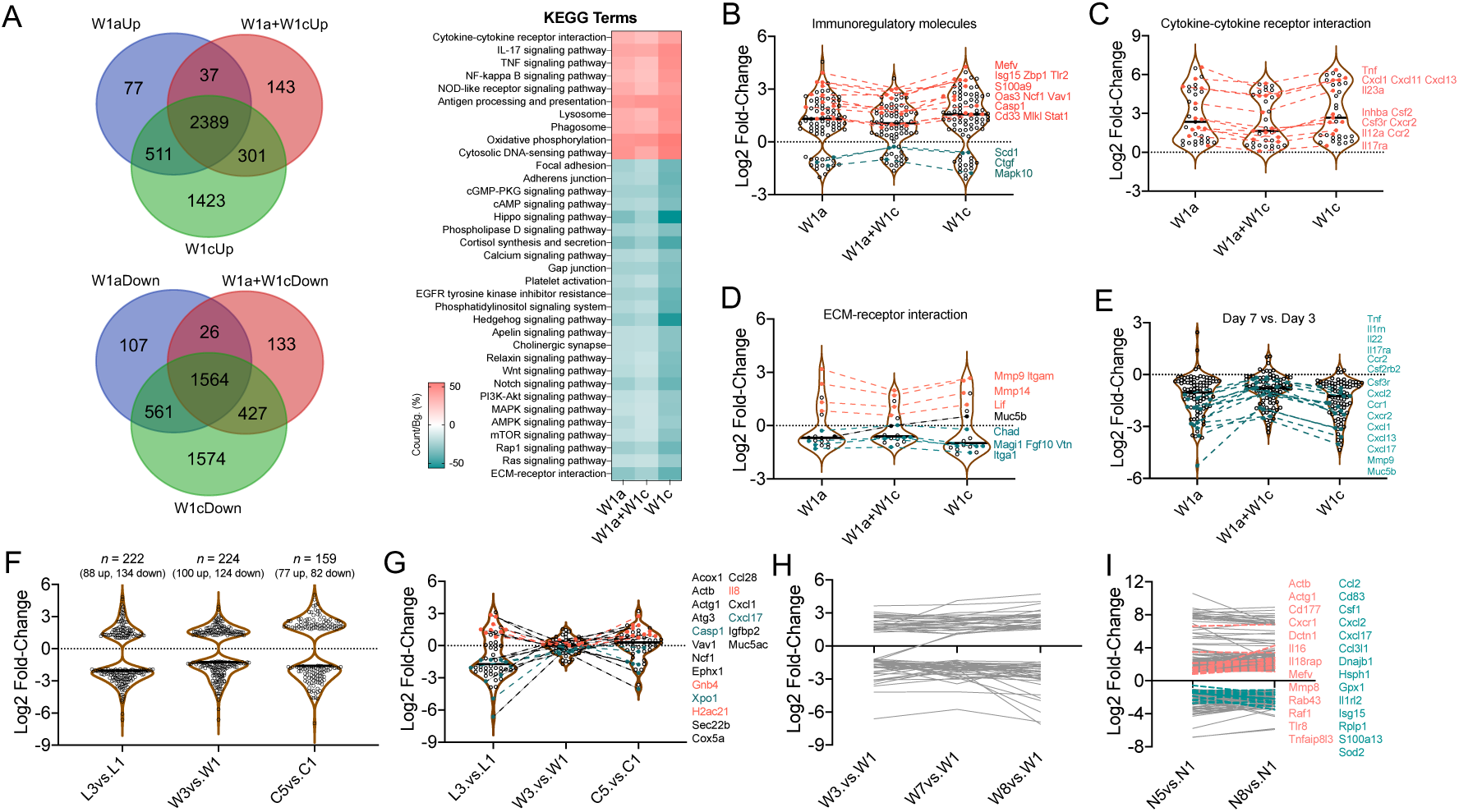
Influence of polymorphic *P. aeruginosa* infection on the inflammatory responses of host lungs. Lung tissues were collected from *P. aeruginosa* W1a and W1c chronically infected mice at day 3 for RNA-seq. (*A*) Venn diagram and KEGG enrichment analysis of the differentially expressed genes. (*B-D*) The significantly changed genes were divided into three categories: (*B*) immunomodulatory molecules, (*C*) cytokine-receptor interaction, and (*D*) ECM-receptor interaction. The fold changes of the differentially expressed genes between the three groups were compared. (*E*) The comparison of the differentially expressed genes between day 3 and day 7. The fold changes of the (*F*) total and (*G*) immune-related differentially expressed proteins between different sampling periods of patient L (the first and the last rounds), patient C (the first and the last rounds), and patient W (rounds 1, 3, 7 and 8). (*H*) The expression levels of immune-related proteins in the BALs from different sampling periods of patient W. (*I*) The expression changes of immune-related genes in the BALs from different sampling periods of patient N.

We then validated the immune responses of the hosts by profiling the proteomes of clinical samples from different sampling periods of patient L (colonized by more QS-intact *P. aeruginosa*), patient C (colonized by QS-deficient *P. aeruginosa*), and patient W (colonized by polymorphic *P. aeruginosa*). The results of mass spectrometry showed that although the samples from round 3 of patients L and W had more significantly expressed proteins compared to the initial sampling period, the immune-related proteins in patient W, who was colonized by polymorphic *P. aeruginosa* population until death showed the smallest expression fluctuation (Figs. 5*F*, *G*, and S14*E*). We further identified that the expression levels of immune-related proteins in the BALs from different sampling periods of patient W remained relatively stable (Fig. 5*H*). We finally evaluated the transcriptional changes of immune-related genes in the BALs from different sampling periods of patient N, who was also colonized by a mixture of QS-intact and QS-deficient *P. aeruginosa*, by using RNA-seq. Indeed, the expression levels of immune-related proteins were less changed during further chronic infection (Fig. 5*I*). Altogether, these results suggested that the genetic evolution of QS mutants during chronic infection in the population greatly influences the immune status of the hosts.

## Discussion

The exertion and evolution of group behaviors in bacterial populations greatly expand the habitats and enhance the survival fitness of these microorganisms. The successful connection of public goods game and bacterial interaction has remarkably facilitated studies focusing on the persistent colonization and pathogenesis of bacteria in the theoretical context of sociomicrobiology (1, 4, 28). In the present study, we characterize the evolutionary dynamics of polymorphic *P. aeruginosa* population in COPD airways and identify a cascaded public goods game mediated by WT cooperators and continuously evolving QS-deficient cheaters that stabilizes social cooperation. Moreover, we find that the involvement of evolved QS-deficient cheaters in *P. aeruginosa* population facilitate the development of chronic infection.

As a star bacterial species in studying the evolution and maintenance of cooperation, *P. aeruginosa* evolves several social traits, especially QS which plays vital roles in nutrient acquisition, defense and virulence of *P. aeruginosa*, and frequently select the mutants deficient in producing the sharable extracellular products (29–33). Previous studies have verified the *de novo* emergence of *lasR* mutant-related QS cheaters among during the *in vitro* evolution of *P. aeruginosa* (17, 34, 35). In the vast majority of cases, QS cheaters would coexist with cooperators in the population and the frequencies of them were dynamically changed. The present study reveals that COPD airways are susceptible to being cocolonized by *P. aeruginosa* with complex pathoadaptive phenotypes and mutations until the end of patients’ lives, typically the mixture of QS-intact and QS-deficient isolates (Fig. 1*C* and Table S1), even though the invasion of some isolates might be attributed to nosocomial interpatient transmission (Fig. 2*A*). This evidence indicates that coexistence of population with polymorphic genetic structure may be an optimal strategy for *P. aeruginosa* to persist in host tissues.

Here we further hypothesize that evolution of polymorphic *P. aeruginosa* population may mainly concentrate on selecting *lasR* mutants that are beneficial to the survival advantage of the population in host tissues. A recent work by Azimi *et al*. (35) studied the long-term evolution of monocultured *P. aeruginosa* PAO1 in synthetic sputum medium. They identified abundant mutant isolates in the early evolving population and a significant positive correlation between *lasR* mutant frequency and the resistance of polymorphic population to β-lactam antibiotics during further evolution. However, it seems that the enhanced antibiotic resistance of the population was simply due to the enrichment of *lasR* mutant, rather than further evolutionary selection on *lasR* mutant that harboring additional resistance-related mutation/s. Because the susceptibility of *lasR* mutant to β-lactam antibiotics could be restored by the complementation of intact *lasR* gene (35). By contrast, by comparing the evolutionary trajectories of monocultured *lasR* mutant and its parental WT strain towards the resistance of ribosome-targeting antibiotics, Hernando-Amado et al. (36) identified a negative epistatic interaction between *lasR* and antibiotic resistance genes by showing that selection on *lasR* mutants and antibiotic resistance are mutually antagonistic. In any case, by characterizing the evolutionary trajectories of polymorphic *P. aeruginosa* population in the lung of COPD patient W, our present study clearly demonstrates that the evolution of the polymorphic population was mainly concentrated on modifying the function of *lasR* mutants (Figs. 1*C* and 2). This can be supported by the successively occurring mutation events in *lasR* mutants in the main group isolates of the same patient. The two subgroups of *P. aeruginosa* isolates were similar to the previously reported CF-adapted *lasR* mutants, which were collected from the later stage of CF airways (27). Typically, the isolate W1c, which might be transmitted from an unknown reservoir and was isolated along with W1a in the present study, as well as all the isolates of patient C, accumulated a large amount of harmful mutations in various pathoadaptive genes and were capable of chronically infecting host lungs without the assistance of QS-intact *P. aeruginosa* (Figs. 2*A*, *B*, 4*A, B*, and Table 1). These results highlight the notion that *P. aeruginosa lasR* mutants may undergo further adaptive genetic evolution during chronic infection and will have a significant competitive advantage over the less evolved individuals.

In the previously reported *in vitro* interaction of cheating on cheaters (26), WT *P. aeruginosa* could be exploited by its isogeneic *lasR* mutant in a QS-controlled product-mediated public goods game, and the *pvdS* mutant could invade the *lasR* mutant in a siderophore-mediated public goods game. Our present study further identifies a cascaded public goods game among the continually evolved *P. aeruginosa* isolates from COPD airways. All the three *lasR* mutants (W1c, W4b, and W8a) could invade W1a in QS-required medium despite their different relative fitness in the competitions, and W8a with an additional mutation in the *pvdS* gene also had a growth advantage over the other isolates in iron-limiting medium (Figs. 3*A-H* and S6), indicative of a phase transition of social interaction in the polymorphic *P. aeruginosa* population. This can be supported by our finding that the emergence of *pvdS* mutation in *P. aeruginosa* isolates from COPD airways was posterior to the *lasR* mutants (Fig. S7). Moreover, cooperators were stabilized in the competitions of W1a, W1c, and W8a, and the frequencies of which differed along with the change of environmental cues (Fig. 3*J-L*). We did not verify the interactions of these clinical isolates by using model *P. aeruginosa* strains with pure genetic backgrounds here, because the additional mutations in W4b and W8a would not influence the functions of QS system and siderophore production. These findings combined with the evolutionary trajectory of *P. aeruginosa* reveal that the QS-controlled extracellular protease-mediated public goods game is the mainstream social interaction within the *P. aeruginosa* population colonizing host lungs. Further evolution towards adjusting the roles of *lasR* mutants will efficiently contribute to stabilizing the population structure according to environmental change, and thus provides an explanation for the persistent colonization and QS-related recurrent attacks of *P. aeruginosa*.

The invasion of *lasR* mutant will reduce the virulence of *P. aeruginosa*, and this might be associated with the decreased production of extracellular proteases at population level (35, 37). Differently, Mould *et al*. (38) identified a cross-feeding interaction between *lasR-*intact and *lasR* mutant *P. aeruginosa* by showing that, the *lasR-*intact individuals would produce citrate upon the stimulation of pyochelin produced by *lasR* mutant, and exclusively activate the *rhl-*QS system of *lasR* mutant to hyperproduce pyocyanin and biofilm. Moreover, LaFayette *et al*. (27) reported that the evolved *lasR* mutants from CF airways could elicit hyperinflammatory responses. These findings combined with the abundant *lasR* variants with altered QS regulation from CF airways (39), collectively imply the functional diversity of *lasR* mutants in polymorphic *P. aeruginosa* population. Our present study investigates the survival advantage of evolved polymorphic *P. aeruginosa* population in host lungs by using *lasR*-intact *P. aeruginosa* and the evolved *lasR* mutant W1c. In contrast to the clearance of *lasR* mutant of *P. aeruginosa* reference strain by the host (40), W1c showed an avirulent phenotype but could successfully colonize mouse lungs (Figs. 4*A*, *B* and S10). Importantly, a mixture of *lasR*-intact *P. aeruginosa* and evolved *lasR* mutant induced the minimum immune response in a mouse model compared to those chronically infected by a single genotype isolate, and a similar trend was also observed in patient samples (Figs. 4 and 5). These data suggest that the interactions of individuals within the evolved polymorphic *P. aeruginosa* population have a significant effect on bacteria-triggered host immune fluctuations or responses during infection. This also indicates that reducing the intensity of the host inflammatory response is crucial for the colonization and long-term survival of pathogenic bacteria. To achieve the above purpose, this may also be a direction of QS mutation evolution. Furthermore, COPD patients have more complex respiratory pathologies than those within CF, and the evolution of multiple independent routes of QS-deficient mutations in COPD can promote the colonization of multiple strains in different infection niches.

Collectively, this study dissects the pathoadaptive evolution and intraspecific interaction of *P. aeruginosa* longitudinally collected from COPD airways. The early evolution of *P. aeruginosa* in lung environments frequently selects mutants deficient in producing the costly and sharable extracellular products, and results in the cocolonization of isolates with diverse QS capacities. We further find that the persistent colonization of the polymorphic *P. aeruginosa* population is associated with the modification of the adaptability of *lasR* mutants, by which a cascaded public goods game mediated by QS-controlled extracellular products and siderophores will be introduced to stabilize the population structure and to promote the population fitness during chronic lung infection. These results demonstrate the multistage evolution and complex interaction of *P. aeruginosa* in adaptation to the host lungs, and provide plausible explanations for the maintenance of cooperation in the game of public goods and the recurrence of *P. aeruginosa*-related chronic infections.

Moreover, the identification of the *lasR* mutant-centered cascaded public goods game in the present study also raises a significant concern regarding the flourishing development of antivirulence drugs targeting a single core regulator of the QS system (41–43). Even the *lasR* mutant may produce the QS-controlled products by invoking the regulation of MvfR (PqsR), a sublevel regulator of the QS system, or elicit other unexpected host immune responses by harboring pathoadaptive mutations during further evolution (44, 45). Therefore, the *in vivo* evolutionary trajectories and interaction dynamics of *P. aeruginosa* isolates identified in the present study, as well as the characterization of host immune responses induced by different combinations of them, provide a vital basis for further understanding the pathogenesis of *P. aeruginosa* and the development of therapeutic strategies against pseudomonal infections.

### Materials and Methods

#### Sample collection, Bacterial strains, and Media

A total of 25 patients (56 to 92 years old) who were diagnosed as COPD and *P. aeruginosa*-positive (but negative in the past whole year) hospitalized in the Affiliated Hospital of Chengdu University were enrolled for longitudinal collection of *P. aeruginosa*. BALs of the patients were recovered with endoscopic surgery using electronic bronchoscope (Olympus BF-10) at an interval of 15 days. Spontaneous sputa in the morning were collected from the patients if the endoscopic surgery was unnecessary. A portion of the sample was cultured in lysogeny broth (LB) at 37 °C with shaking (220 rpm) for two days. The culture liquid was spread on LB plate and cultured at 37 °C overnight. The colonies with apparent differences in shape, color, size and surface states were picked out for 16S rDNA-based species identification. The single colony of identified *P. aeruginosa* clinical isolates and the reference isolate *P. aeruginosa* PAO1 were cultured in LB medium and preserved at – 80 °C for further use.

#### Ethical Statement

BALs and sputum samples were obtained from the COPD patients hospitalized in the affiliated hospital of Chengdu University (Chengdu, China). Written informed consents were received from the patients or their immediate family members. The study was approved by the Ethics Committee of the Affiliated Hospital of Chengdu University (PJ2020-021-03), and all methods were carried out in accordance with the guidelines and regulations of Chengdu University. Mouse models used in this study were bought from Beijing Huafukang Laboratory Animal Co., Ltd. (Beijing, China) and housed in the specific-pathogen-free facility of the State Key Laboratory of Biotherapy, Sichuan University. Animal experiments were approved by the Ethics Committee of the State Key Laboratory of Biotherapy (2021559A) and carried out in compliance with institutional guidelines concerning animal use and care of Sichuan University.

#### Isolate typing and phenotypic identification

Genomic DNAs of overnight cultured *P. aeruginosa* isolates were harvested using Bacterial DNA Isolation Kit (Foregene Biotechnology, Co. Ltd., China). Typing of *P. aeruginosa* isolates was performed by ERIC-PCR using the single primer 5’- AACTAAGTAACTGGGGTGAGCG-3’ (46). Phenotypic identification of *P. aeruginosa* isolates was performed as recommended by Filloux and Ramos (47) using PAO1 as positive control. In brief, *P. aeruginosa* was inoculated on M9-adenosine (0.1%, w/v) or M9-skim-milk (0.5%, w/v) plates to test the production of QS-controlled intracellular protease Nuh or extracellular proteases by checking the growth or the size of proteolytic halo, respectively. Pyocyanin production was determined by extracting pyocyanin from the supernatant using chloroform and HCl, followed by measuring the absorbance at 520 nm. Biofilm production was determined by crystal violet staining followed by measuring the absorbance at 590 nm. Swimming and twitching motilities were determined by measuring the diameters of colonies cultured on M8 plates containing 0.3% or 1.0% agar powder. The production of siderophores was determined by measuring the cell densities of *P. aeruginosa* cultured in M9-CAA medium supplemented with 50 μM of FeCl_3_ for 24 h. All the experiments were independently repeated for three times. The phenotypic values of *P. aeruginosa* clinical were compared to those of PAO1.

#### WGS, genomic and phylogenetic analyses

Libraries of *P. aeruginosa* genomic DNAs were constructed by using NEBNext^®^Ultra™ DNA Library Prep Kit for Illumina (New England Biolabs, USA), and then WGS was performed on the Illumina HiSeq PE150 platform (Novogene Bioinformatics Technology Co. Ltd., China). The raw data are deposited in the NCBI BioProject database under accession number PRJNA846307. High quality pair-end reads were assembled by using the software SOAP denovo v2.04, SPAdes, and ABySS, and integrated by CISA, followed by sequence optimization using Gapclose v1.12. The program GeneMarkS was used to retrieve the coding sequences, and the gene functions were predicted by using the databases of GO, KEGG, COG, NR, TCDB and Swiss-Prot. MUMmer and LASTZ were used to identify the SNPs and InDels in the genomes of *P. aeruginosa* clinical isolates from the later sampling periods compared to those of the corresponding isolates from the initial. SnpEff v4.3 was used to evaluate the mutation impact of SNPs. The common and different numbers of SNPs, InDels and genes were sorted by using Venn diagram (http://bioinformatics.psb.ugent.be/webtools/Venn/). The assembled contigs of *P. aeruginosa* sequenced in this study and the complete genome sequences of *P. aeruginosa* PAO1 (NCBI accession no. AE004091.2), PA14 (NCBI accession no. NC_008463.1), PAK (NCBI accession no. CP020659.1), and DK2 (NCBI accession no. NC_018080.1) were used to construct SNP-based phylogenetic tree with kSNP v3.0. The genome sequence of *Azotobacter vinelandii* (NCBI accession no. NC_012560.1) was set as the outgroup. The tree was visualized with FigTree v1.4.3 (http://tree.bio.ed.ac.uk/software/figtree/). Genome sequence-based typing of MLST and serotype were performed by using the online software MLST v2.0 (https://cge.food.dtu.dk/services/MLST/) and PAst v1.0 (https://cge.food.dtu.dk/services/PAst/).

#### Transcriptomic analysis of *P. aeruginosa* isolates

The total RNAs of *P. aeruginosa* isolates (cultured to cell densities of OD_600_ = 1.5 in LB broth) from different sampling periods of COPD patients were isolated using Total RNA Isolation Kit with gDNA removal (Foregene Biotechnology, Co. Ltd., China). Each RNA sample from three independent biological replicates were mixed. RNA samples from two parallel experiments were conducted for library construction using NEBNext^®^Ultra™ RNA Library Prep Kit for Illumina, followed by prokaryotic strand-specific RNA-seq on the Illumina HiSeq PE150 platform (Novogene Bioinformatics Technology Co. Ltd., China). The raw data are deposited in the NCBI BioProject database under accession number PRJNA846307. High quality pair-end reads were mapped to the genome of *P. aeruginosa* PAO1 by Bowtie 2 v2.2.3. Differential gene expression was calculated by HTSeq v0.6.1 and DESeq 2 using expected fragments per kilobase of transcript per million fragments (FPKM). Combined application of KOBAS v2.0, GOseq R package and DAVID v6.8 were used to get the functional categories of KEGG pathway and GO enriched by the differentially expressed genes. The common and different numbers of differentially expressed genes, the enriched KEGG and GO terms among isolates were sorted by using Venn diagram.

#### Competition assay

The interactions of *P. aeruginosa* clinical isolates W1a (*lasR*-intact), W4b (*lasR* mutant), W8a (*lasR-pvdS* mutant), and W1c (evolved *lasR* mutant) were studied by coculturing different combinations of them under different conditions. For LasR-controlled elastase-related two-player competition assay, W1a and W1c, W1a and W4b, W1a and W8a, or W1c and W4b were cocultured in 2 mL of M9-casein medium from different initial ratios (99:1, 1:1, and 99:1) for 24 h with shaking. For PvdS-controlled siderophore-related two-player competition assay, W4b and W8a, W1a and W1c, W1a and W8a, or W1c and W8a were cocultured in 2 mL of M9-CAA medium supplemented with Transferrin (100 μg/mL) from an initial ratio of 99:1 for 24 h with shaking. For the three-player competition assay, a mixture of W1a, W1c, and W8a at the ratio of 98:1:1 was inoculated in 2 mL of M9-casein medium or the medium supplemented with 50 μM of FeCl_3_ or 100 μg/mL of Transferrin, and then successively subcultured (1:10 dilution) in accordant fresh medium at 24 h interval. The experiment was stopped when the frequency of each isolate in the culture was relatively stable. All the experiments were independently repeated for six times. The total cell density of each culture was measured by CFU enumeration on LB plates, and the proportion of each isolate in the coculture was determined by spreading the same volume of culture liquids on different selection plates. Specifically, among the four tested *P. aeruginosa* clinical isolates, only W1a produced a large and apparent proteolytic ring on M9-casein plate, and only W8a showed an Imipenem resistance and could be selected on LB-Imipenem (5 μg/mL) plate. The relative fitness (*v*) of one kind of isolates was calculated using the equation *v* = log_10_[x_1_(1 − x_0_)/ x_0_(1 − x_1_)], where x_0_ indicates the initial frequency and x_1_ indicates the final, and *v* > 0 indicates one isolate grows faster than the opponent, while v < 0 indicates an opposite growth status, as described elsewhere (17).

#### Caenorhabditis elegans Killing Assay

The pathogenicity of *P. aeruginosa* isolates W1a, W1c, and mixture (1:1) of them were determined by using *C. elegans* slow-killing assay as described previously (23). In brief, each *P. aeruginosa* solution was spread on nematode growth medium and incubated at 37 °C for 24 h. Subsequently, 15 newly cultured nematodes at L4 stage were seeded on each plate and further incubated at 25 °C. The survival of nematodes was recorded at 24 h interval. Nematodes fed with *Escherichia coli* OP50 (uracil auxotrophy) were set as control.

#### Mouse Models

Agar-beads-embedded *P. aeruginosa* isolates W1a, W1c, and mixture (1:1) of them, as well as a mixture of W1a, W1c, and W8a (98:1:1) were prepared and used to chronically infect the lung of C57BL/6 mouse (8-week-old, female) as previously described (31). The agar beads were dispersed into 1.0–2.0 × 10^6^ CFUs in 50 μL of sterile saline and intranasally instilled into anaesthetized mice. The survival of mice was recorded at 12 h interval. Three randomly selected mice were killed at designated sampling points. The whole lungs were aseptically removed, homogenized, and conducted for CFU enumeration, RNA isolation and flow cytometry.

#### Flow cytometry

The lung tissues were homogenized in RPMI-1640 and 10% FBS containing 0.2 mg/ml collagenase type I/IV and single-cell suspensions were generated from lung tissues using a gentle MACS™ Octo Dissociator with Heaters following the manufacturer’s protocol (Miltenyi Biotec). The digested lung tissues were filtered through 70 μM cell strainers and red blood cells were lysed using red blood cell lysis buffer (Beyotime, C3702). The resulting single-cell suspensions were washed with PBS and resuspended in PBS containing 1% FBS. The cell suspensions were incubated with FcR Blocking Reagent (Biolegend, catalog no. 101319) for 15 min at 4°C, stained with fluorescent conjugated Abs for 30 min at 4°C and washed twice with PBS. Positive staining with the BD fixable viability dye FVS620 (BD Biosciences, catalog no. 564996) was used to exclude dead cells. Data were collected on a BD FACSymphony analyzer (BD Biosciences) and analyzed with FlowJo software (v.10.4).

#### Quantitative Protein Mass Spectrometry

The proteins in the BALs of patients L and W, and in the sputum samples of patient C were collected and conducted for label-free LC-MS/MS using Orbitrap Exploris 480 matched with FAIMS (Thermo Fisher) with ion source of Nanospray FlexTM (ESI). All the resulting spectra were separately searched against UniProt database for *Homo sapiens* and *P. aeruginosa* by Proteome Discoverer 2.4 (Thermo). The identified peptide spectrum matches and proteins with FDR no more than 1.0% were retained. The protein quantitation results were statistically analyzed by *t*-test. GO and InterPro functional analysis were conducted using the Interproscan program against the non-redundant protein database (including Pfam, PRINTS, ProDom, SMART, ProSite, PANTHER), and the databases of COG (Clusters of Orthologous Groups) and KEGG were used to analyze the protein family and pathway. The common and different numbers of differentially expressed proteins, the enriched KEGG and GO terms among isolates were sorted by using Venn diagram.

#### Transcriptomic analysis of host lungs

The total RNAs of mouse lungs and BALs of COPD patient N were isolated, sequenced, and analyzed as described in ‘Transcriptomic analysis of *P. aeruginosa* isolates’ in the Methods section. Differently, the obtained pair-end reads were mapped to the genome of *Mus musculus* or *Homo sapiens* before the step of differential expression analysis.

#### Statistical Analysis

Graphpad Prism v8.0 (San Diego, CA, USA) was used to process the data generated by the phenotypic identification assays. Mean values of standard deviation were compared by using two-tailed unpaired *t*-test or One-Way ANOVA. The survival curves of *C. elegans* and mice were compared by using Log-rank (Mantel-Cox) test.

## Acknowledgments

This work was supported by the National Natural Science Foundation of China (32270121, 31970131, 81922042 and 82172285), the Sichuan Science and Technology Program (2021JDJQ0042), the high-level talent training program of Chengdu University (2081920066), the 1·3·5 project of excellent development of discipline of West China Hospital of Sichuan University (ZYYC21001), and the innovation foundation of the Affiliated Hospital of Chengdu University (CDFYCX202209).

## Author contributions

K.Z. and X.Z. designed, initiated, and conceived the project. K.Z., X.Y., Yi.Z., He.L., J.S.L., Hu.L., Y.W. and T.H. performed the experiments. K.Z., Q.Z., G.H., Ya.Z. and H.Z. coordinated clinical sample collection. K.Z., L.D. and X.Z. performed computational analyses. K.Z., X.Y., He.L., J.S.L., and X.Z. assisted with mouse experiments and prepared mouse samples. K.Z., X.Y. and Y.C. assisted with the nematode infection experiments. X.W. and Y.C. provided critical reagents and equipment. K.Z. and X.Z. wrote the manuscript with extensive input from all authors. All authors have read and approved the final manuscript.

## Supplementary Figures

**Figure S1.**
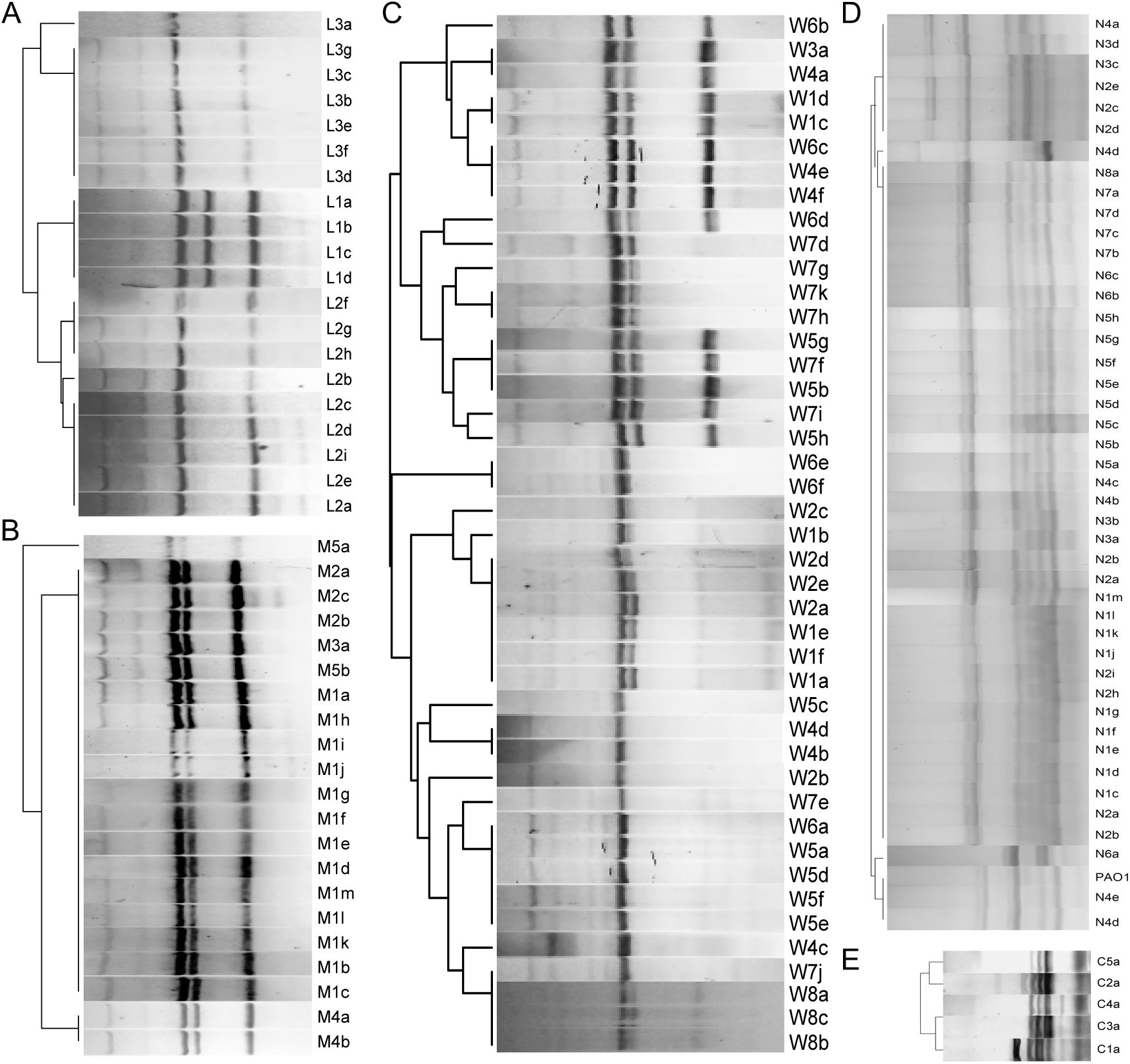
ERIC-PCR-based typing of longitudinally collected *P. aeruginosa* COPD isolates from patients (*A*) L, (*B*) M, (*C*) W, (*D*) N, and (*E*) C.

**Figure S2.**
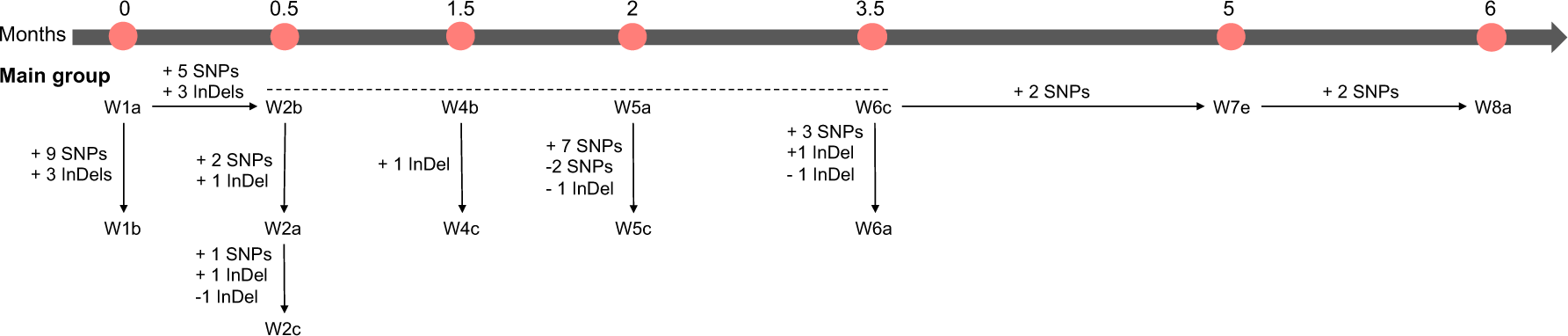
Time-dependent delivery of mutations among the isolates in the main group of *P. aeruginosa* isolates in patient W.

**Figure S3.**
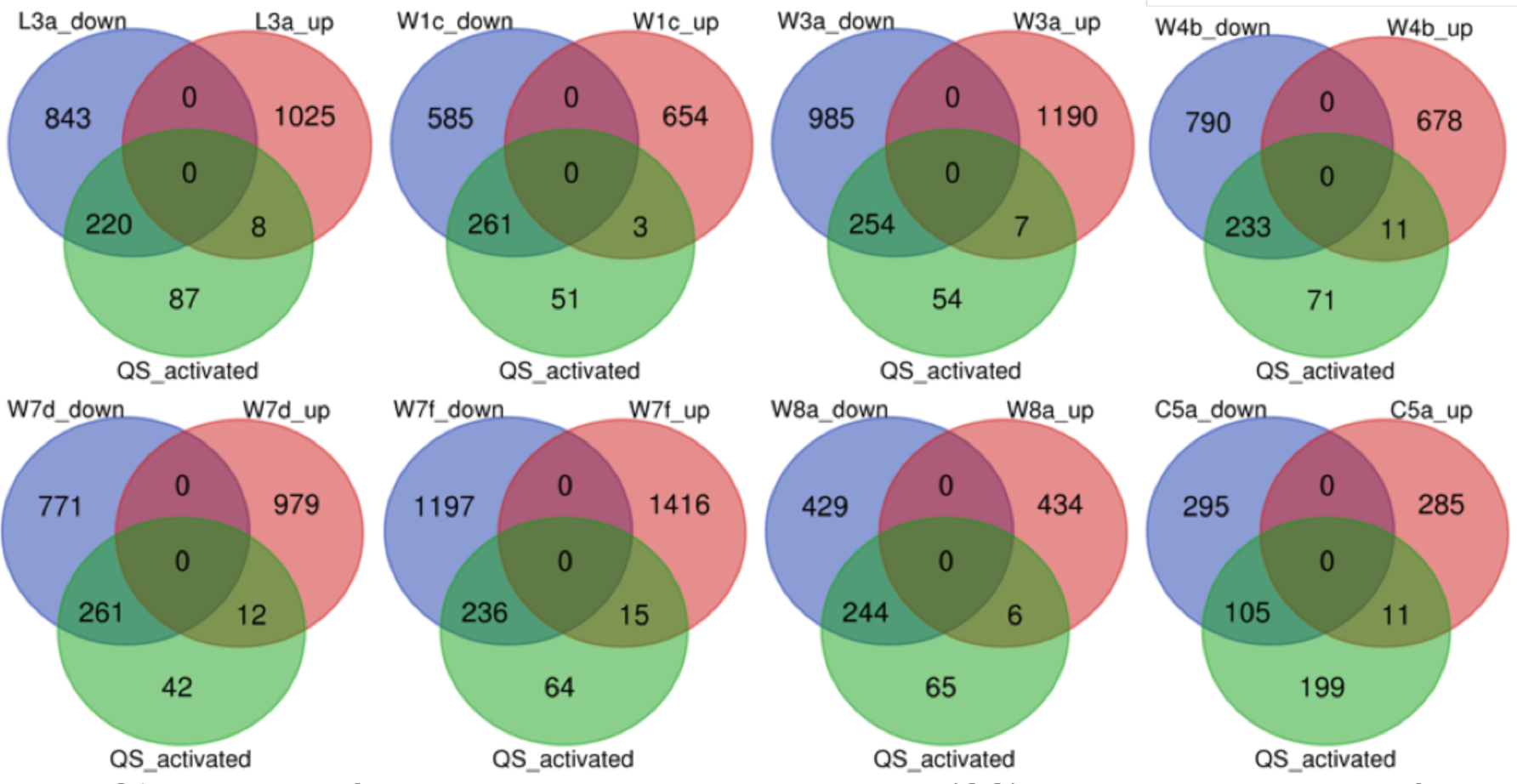
Numbers of genes activated by quorum-sensing (QS) system among the significantly down-and up-regulated genes of evolved *P. aeruginosa* COPD isolates compared to their corresponding initial isolates (*P* < 0.05).

**Figure S4.**
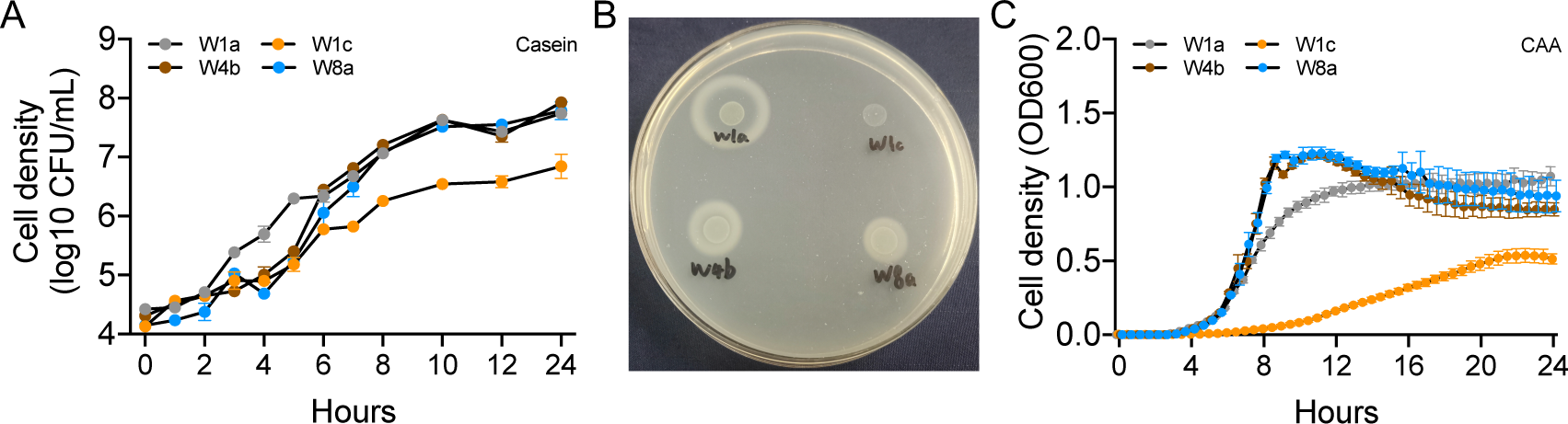
Growth of successively identified *P. aeruginosa* isolates in (*A*) M9-casein (0.5%) broth, on (*B*) M9-casein (0.5%) plate or in (*C*) M9-CAA (0.1%) broth. Data shown are means ± standard deviation (SD) of (*A*) four and (*B*) eight independent replicates.

**Figure S5.**
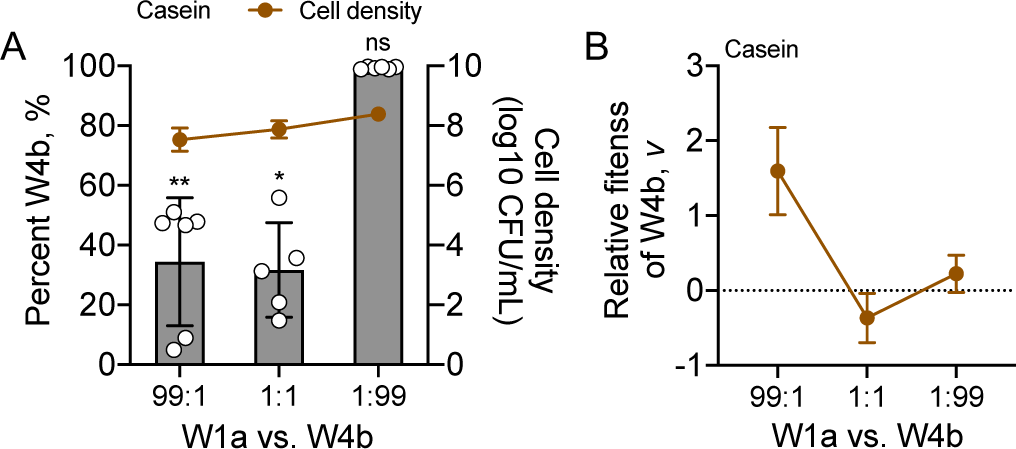
*In vitro* competition of *P. aeruginosa* W1a and W4b in M9-casein medium. (*A*) Frequency and (*B*) relative fitness of W4b in the coculture for 24 h. Data shown are means ± SD of six independent replicates. The value of each column was compared to the initial frequency of corresponding isolate using two tailed unpaired *t-*test. *, *P* < 0.05. **, *P* < 0.01. ns, not significant.

**Figure S6.**
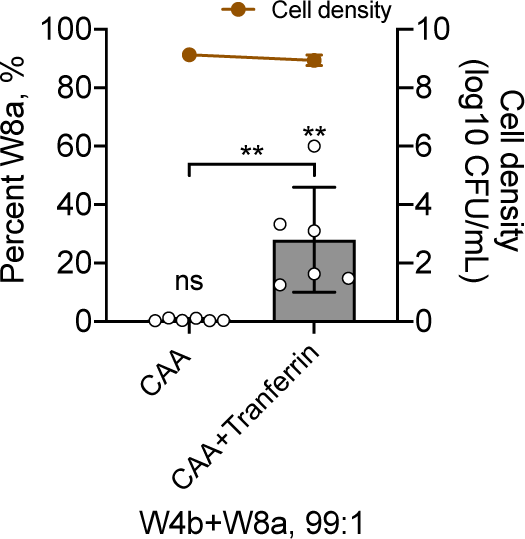
*In vitro* competition of *P. aeruginosa* W4b and W8a in M9-CAA medium with or without iron-limiting. Right Y-axis indicates the cell densities of each culture. Data shown are means ± SD of six independent replicates. The value of each column was compared to the initial frequency of corresponding isolate, or between the two groups using two tailed unpaired *t-*test. **, *P* < 0.01. ns, not significant.

**Figure S7.**
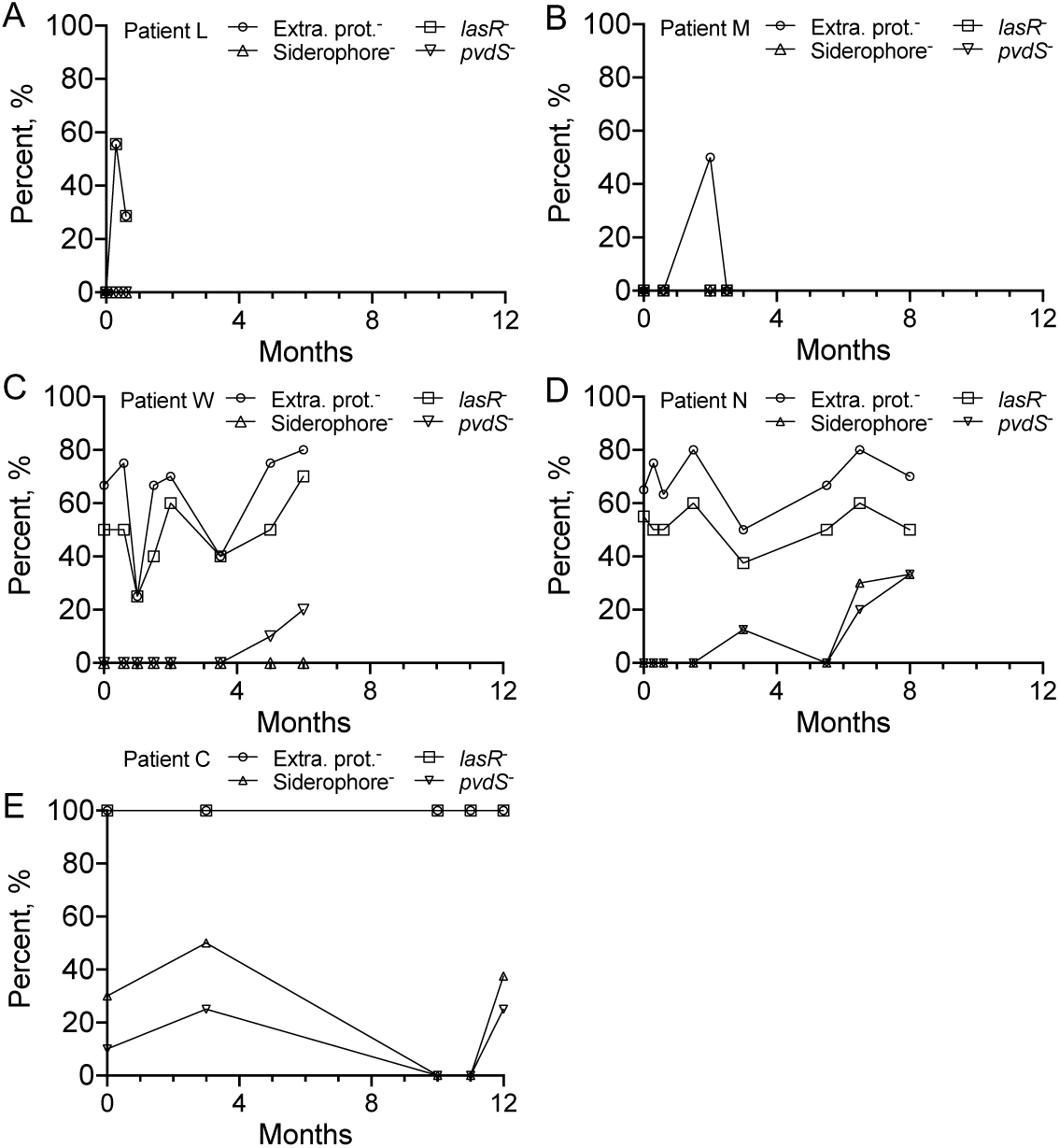
Capacities of *P. aeruginosa* COPD isolates in producing QS-controlled extracellular proteases and siderophores as sampling time increased. Extra. prot.^−^, isolate deficient in producing extracellular protease. *lasR*^−^, *lasR* mutant. siderophore^−^, isolate deficient in producing siderophores. *pvdS*^−^, *pvdS* mutant.

**Figure S8.**
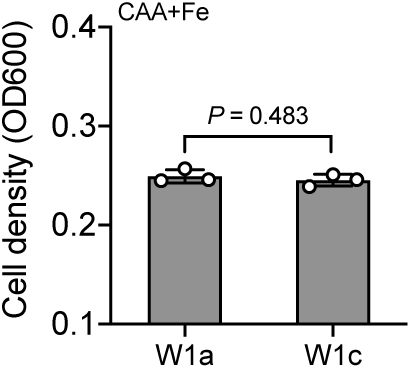
Cell densities of monocultured *P. aeruginosa* W1a and W1c in M9-CAA medium containing 50 μM of FeCl_3_ for 24 h. Data shown are means ± SD of three independent replicates. Two tailed unpaired *t-*test.

**Figure S9.**
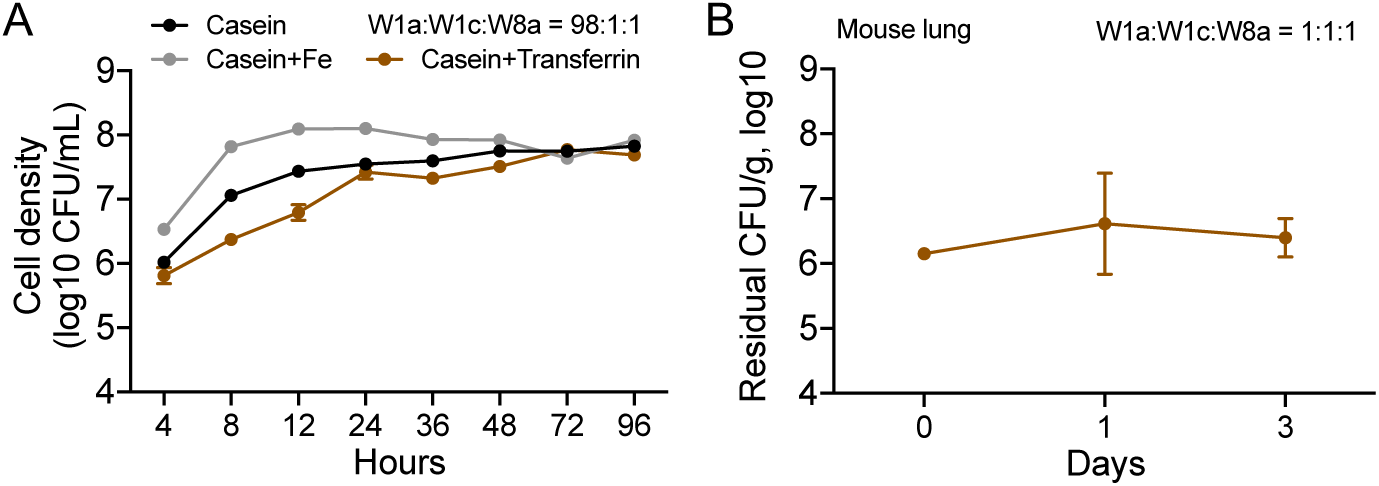
Growth curves of *P. aeruginosa* W1a, W1c, and W8a cocultured under different conditions. (*A*) Cell densities of W1a, W1c, and W8a sub-cocultured in different media from an initial ratio of 98:1:1. The culture media were refreshed at 24 h interval and the experiment were stopped when the frequency of each isolate in the culture was relatively stable. Data shown are means ± SD of six independent replicates. (*B*) Residual colony forming units (CFU) in mouse lungs coinfected by W1a, W1c, and W8a from an initial ratio of 98:1:1. Data shown are means ± SD of five independent replicates.

**Figure S10.**
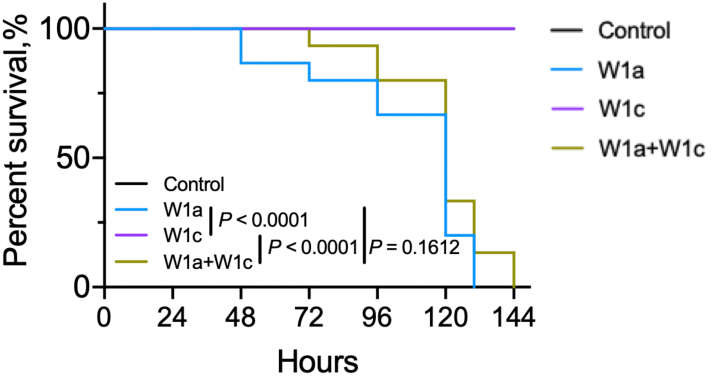
Slow-killing of *Caenorhabditis elegans* infection model (15 nematodes per group) using *P. aeruginosa* isolates W1a, W1c, and 1:1 mixture of them. The survival curves of *C. elegans* and were compared by using Log-rank (Mantel-Cox) test.

**Figure S11.**
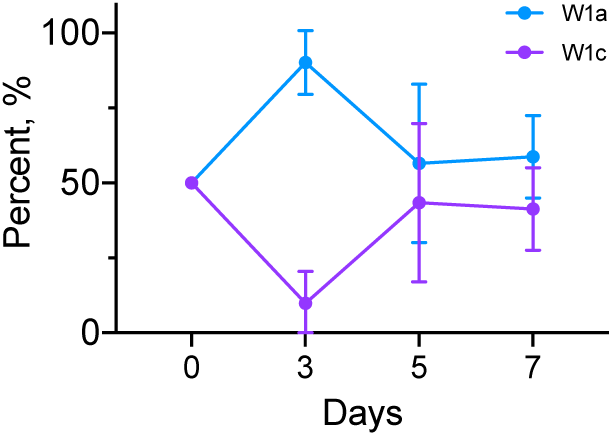
Proportion changes of *P. aeruginosa* isolates W1a and W1c in the lungs of mice chronically infected by 1:1 mixture of W1a and W1c. Data shown are means ± SD of three independent replicates.

**Figure S12.**
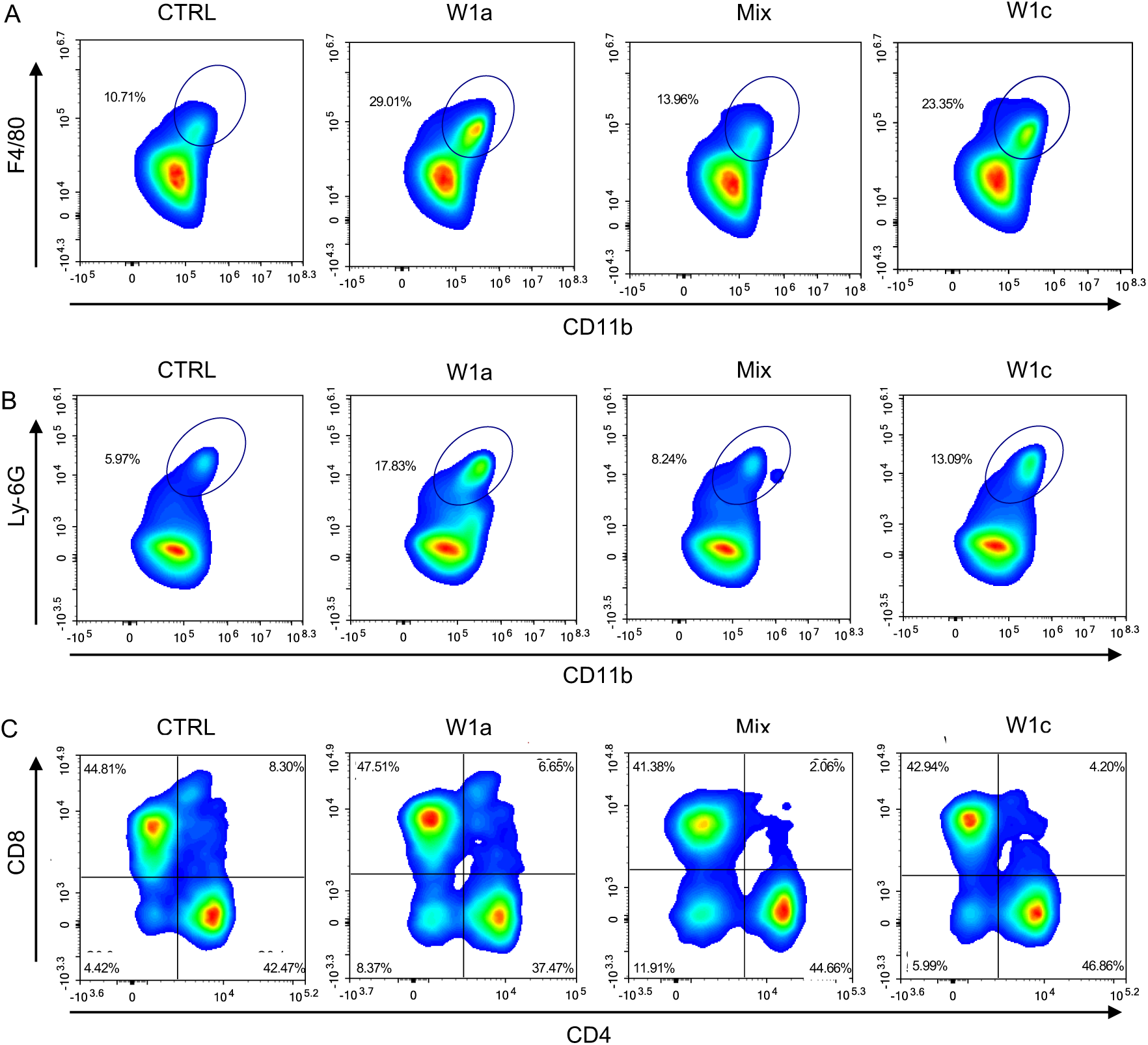
Representative flow cytometry plots of immune cells at day 3. *P. aeruginosa* COPD isolates-infected lung tissues were collected and the proportional changes in inflammatory (*A*) macrophages, (*B*) neutrophils, (*C*) CD8^+^ T cells and CD4^+^ T cells were detected under the gate of CD45^+^ in the different groups.

**Figure S13.**
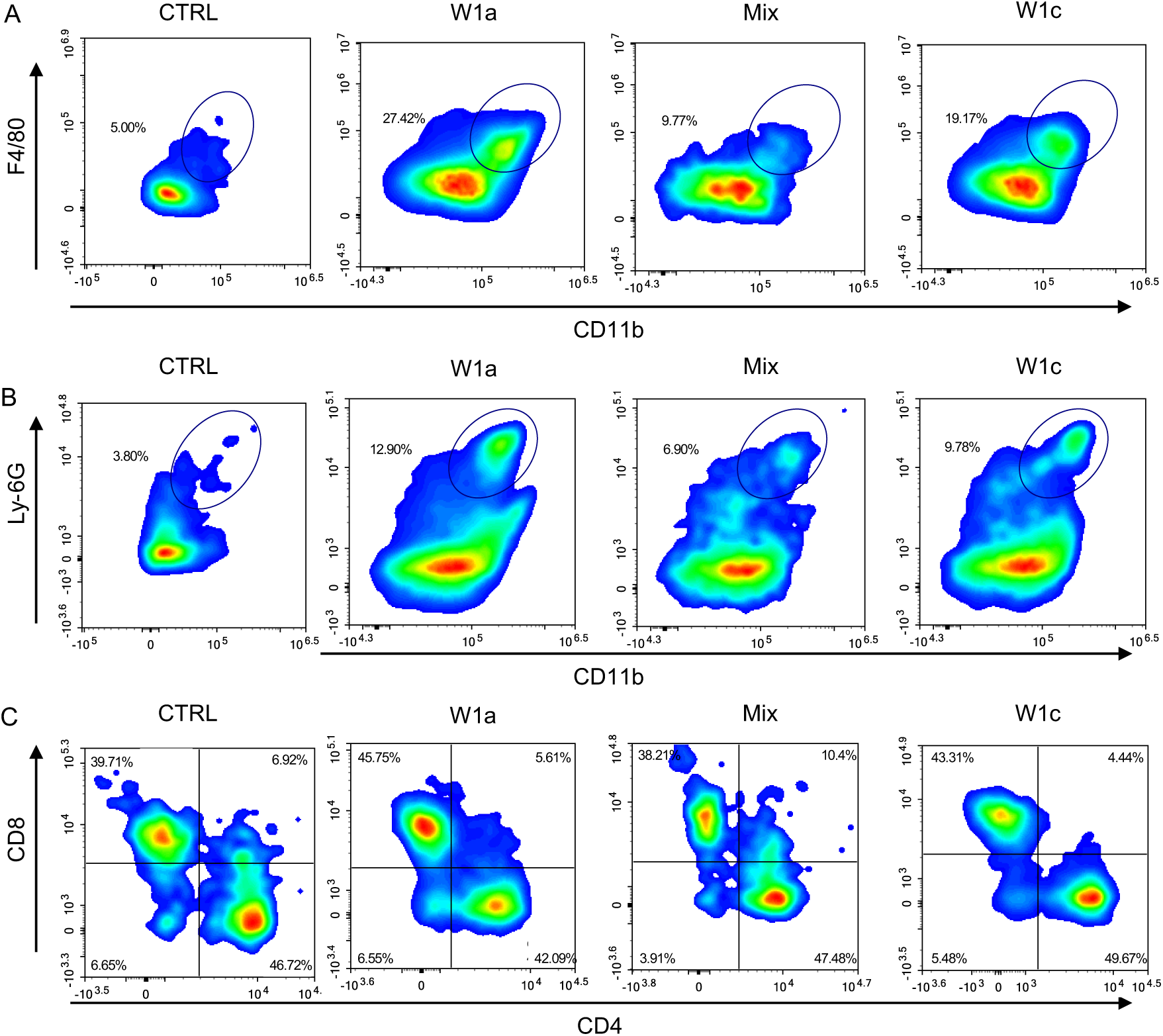
Representative flow cytometry plots of immune cells at day 7. *P. aeruginosa* COPD isolates-infected lung tissues were collected and the proportional changes in inflammatory (*A*) macrophages, (*B*) neutrophils, (*C*) CD8^+^ T cells and CD4^+^ T cells were detected under the gate of CD45^+^ in the different groups.

**Figure S14.**
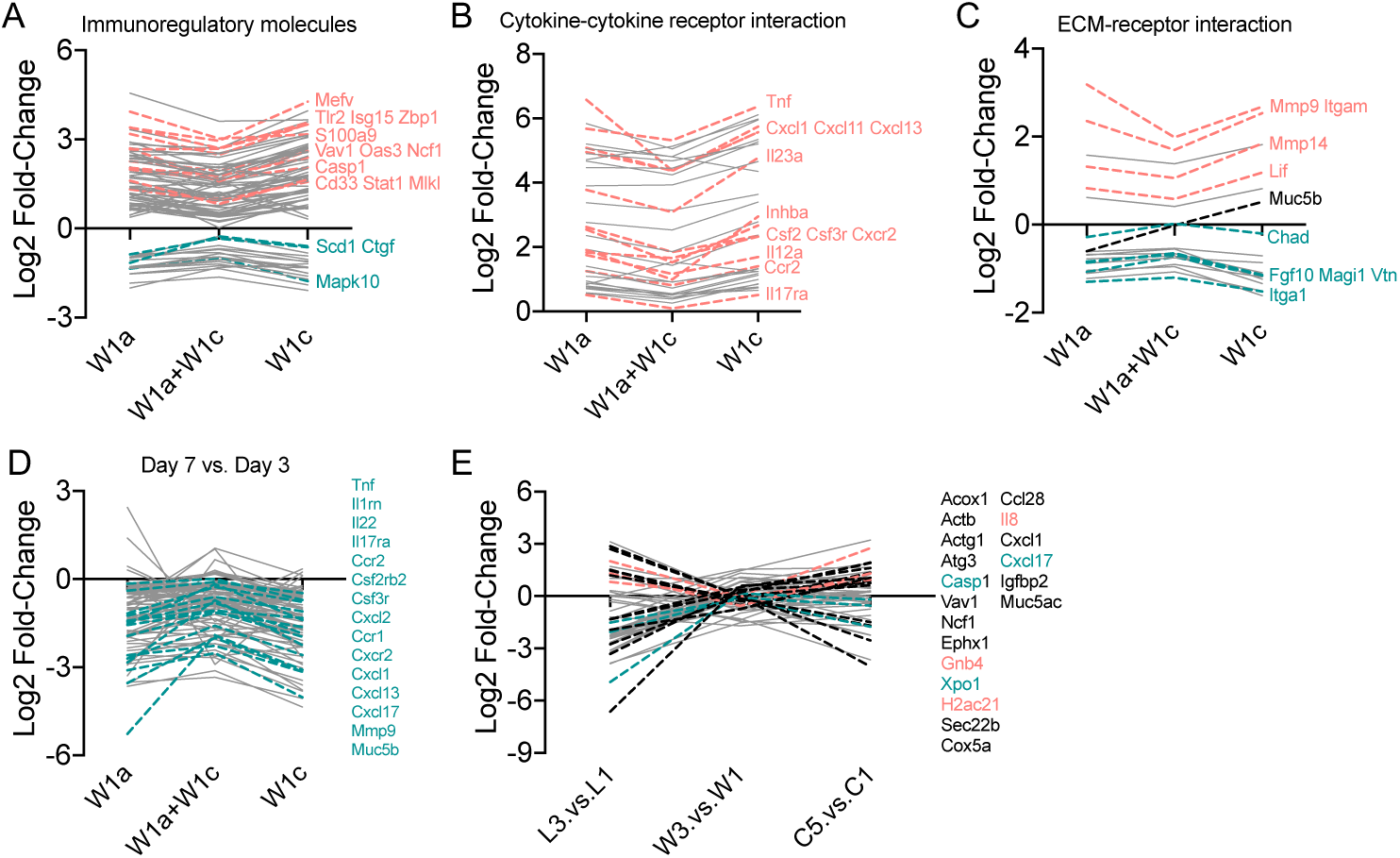
The effects of the interaction of QS-intact and QS-deficient *P. aeruginosa* on differentially expressed genes (different data presentation, related to Figure 5). (*A-C*) The fold changes of the differentially expressed genes from *P. aeruginosa* W1a and W1c chronically infected mouse lung tissues at day 3. The immune-related genes were sub-clustered into (*A*) immunomodulatory molecules, (*B*) cytokine-receptor interaction and (*C*) ECM-receptor interaction. (*D*) The changes fold of the differentially expressed genes between day 3 and day 7. (*E*) The fold changes of the immune-related differentially expressed proteins between different sampling periods of patient L (the first and the last rounds), patient C (the first and the last rounds), and patient W (rounds 1, 3, 7 and 8).

## Supplementary Tables

**Table S1.** Information of whole-genome sequencing of P. aeruginosa COPD isolates.

| Sample name | Scaffold Num (>500bp) | Total Length (bp) | N50 Length (bp) | N90 Length (bp) | Max Length (bp) | Min Length (bp) | GC Content (%) | Gene Num |
| --- | --- | --- | --- | --- | --- | --- | --- | --- |
| L1a | 48 | 6,194,072 | 427,176 | 101,614 | 582,926 | 602 | 66.54 | 5,639 |
| L2a | 28 | 6,435,842 | 427,034 | 139,968 | 927,411 | 1,126 | 66.38 | 5,942 |
| L2b | 28 | 6,444,043 | 440,552 | 140,180 | 927,971 | 1,255 | 66.38 | 5,949 |
| L3a | 30 | 6,441,852 | 440,553 | 139,968 | 927,655 | 993 | 66.38 | 5,947 |
| L3b | 27 | 6,434,508 | 440,553 | 140,093 | 927,440 | 1,255 | 66.38 | 5,943 |
| M1a | 25 | 6,455,667 | 773,339 | 181,302 | 926,780 | 1,255 | 66.46 | 5,955 |
| M2a | 32 | 6,507,772 | 484,419 | 106,705 | 876,866 | 754 | 66.47 | 6,006 |
| M3a | 16 | 6,410,450 | 773,326 | 216,382 | 926,780 | 63,072 | 66.48 | 5,929 |
| M4a | 16 | 6,343,091 | 538,911 | 216,391 | 868,624 | 63,101 | 66.48 | 5,864 |
| M5a | 29 | 6,459,656 | 773,326 | 181,290 | 876,866 | 1,174 | 66.46 | 5,958 |
| W1a | 52 | 6,552,282 | 289,459 | 93,048 | 670,307 | 1,051 | 66.3 | 6,016 |
| W1b | 32 | 6,541,402 | 407,011 | 115,986 | 669,860 | 7,831 | 66.31 | 6,062 |
| W1c | 26 | 6,373,967 | 665,413 | 173,676 | 930,301 | 1,096 | 66.35 | 5,868 |
| W2a | 52 | 6,587,294 | 288,721 | 93,047 | 669,158 | 629 | 66.28 | 6,072 |
| W2b | 50 | 6,512,382 | 289,505 | 116,034 | 670,319 | 1,051 | 66.32 | 5,971 |
| W2c | 49 | 6,624,046 | 407,011 | 93,048 | 669,158 | 629 | 66.27 | 6,126 |
| W3a | 25 | 6,456,802 | 773,326 | 216,382 | 926,780 | 1,174 | 66.46 | 5,961 |
| W4a | 28 | 6,373,960 | 665,413 | 117,991 | 930,301 | 1,096 | 66.35 | 5,869 |
| W4b | 52 | 6,633,806 | 293,890 | 115,980 | 670,070 | 611 | 66.26 | 6,133 |
| W4c | 49 | 6,579,997 | 407,011 | 93,155 | 670,307 | 1,051 | 66.33 | 6,018 |
| W5a | 51 | 6,718,523 | 270,086 | 115,915 | 669,083 | 1,051 | 66.32 | 6,180 |
| W5b | 25 | 6,457,074 | 773,318 | 181,302 | 926,780 | 1,255 | 66.46 | 5,964 |
| W5c | 48 | 6,623,350 | 270,086 | 93,048 | 892,561 | 1,115 | 66.27 | 6,121 |
| W6a | 55 | 6,742,645 | 288,667 | 93,245 | 708,299 | 629 | 66.23 | 6,245 |
| W6b | 28 | 6,374,455 | 665,413 | 173,786 | 904,071 | 1,096 | 66.35 | 5,867 |
| W6c | 50 | 6,625,247 | 294,441 | 93,246 | 670,094 | 629 | 66.27 | 6,128 |
| W7d | 39 | 6,463,062 | 426,565 | 110,820 | 727,067 | 591 | 66.49 | 5,866 |
| W7e | 51 | 6,530,340 | 307,344 | 89,431 | 670,094 | 629 | 66.32 | 5,975 |
| W7f | 24 | 6,499,873 | 773,326 | 216,382 | 926,780 | 754 | 66.46 | 5,992 |
| W8a | 50 | 6,512,839 | 288,721 | 90,245 | 670,319 | 1,051 | 66.32 | 5,971 |
| C1a | 29 | 6,466,799 | 471,277 | 142,054 | 876,866 | 1,174 | 66.46 | 5,963 |
| C5a | 37 | 6,322,317 | 437,793 | 134,783 | 965,968 | 1,128 | 66.45 | 5,756 |

**Table S2.** Summary of InDel (insertion and deletion) sites in the genomes of evolved *P. aeruginosa* COPD isolates compared to corresponding initial isolates.

| Sample name | Total INDEL | CDS INDEL | Intergenic INDEL | Start codon ins | Start codon del | CDS inside del | Stop codon ins | Stop codon del | CDS inside ins | Frame-shifted INDEL |
| --- | --- | --- | --- | --- | --- | --- | --- | --- | --- | --- |
| W1b | 8 | 3 | 5 | 0 | 0 | 1 | 0 | 0 | 2 | 0 |
| W1c | 360 | 51 | 309 | 1 | 2 | 23 | 1 | 0 | 24 | 16 |
| W2a | 10 | 4 | 6 | 0 | 0 | 2 | 0 | 0 | 2 | 0 |
| W2b | 9 | 3 | 6 | 0 | 0 | 1 | 0 | 0 | 2 | 0 |
| W2c | 8 | 4 | 4 | 0 | 0 | 3 | 0 | 0 | 1 | 1 |
| W3a | 362 | 50 | 312 | 2 | 2 | 28 | 1 | 0 | 17 | 9 |
| W4a | 361 | 50 | 311 | 1 | 2 | 23 | 1 | 0 | 23 | 16 |
| W4b | 10 | 4 | 6 | 0 | 0 | 2 | 0 | 0 | 2 | 0 |
| W4c | 11 | 4 | 7 | 0 | 0 | 2 | 0 | 0 | 2 | 1 |
| W5a | 8 | 3 | 5 | 0 | 0 | 1 | 0 | 0 | 2 | 0 |
| W5b | 366 | 51 | 315 | 2 | 2 | 29 | 1 | 0 | 17 | 10 |
| W5c | 7 | 3 | 4 | 0 | 0 | 2 | 0 | 0 | 1 | 0 |
| W6a | 7 | 3 | 4 | 0 | 0 | 1 | 0 | 0 | 2 | 0 |
| W6b | 364 | 52 | 312 | 1 | 2 | 24 | 1 | 0 | 24 | 17 |
| W6c | 10 | 4 | 6 | 0 | 0 | 2 | 0 | 0 | 2 | 0 |
| W7d | 388 | 61 | 327 | 2 | 1 | 29 | 1 | 0 | 28 | 16 |
| W7e | 9 | 3 | 6 | 0 | 0 | 1 | 0 | 0 | 2 | 0 |
| W7f | 364 | 51 | 313 | 2 | 2 | 29 | 1 | 0 | 17 | 10 |
| W8a | 10 | 3 | 7 | 0 | 0 | 1 | 0 | 0 | 2 | 0 |
| M2a | 1 | 0 | 1 | 0 | 0 | 0 | 0 | 0 | 0 | 0 |
| M3a | 1 | 0 | 1 | 0 | 0 | 0 | 0 | 0 | 0 | 0 |
| M4a | 1 | 0 | 1 | 0 | 0 | 0 | 0 | 0 | 0 | 0 |
| M5a | 2 | 1 | 1 | 0 | 0 | 0 | 0 | 0 | 1 | 1 |
| L2a | 34 | 5 | 29 | 0 | 0 | 3 | 0 | 0 | 2 | 1 |
| L2b | 34 | 5 | 29 | 0 | 0 | 3 | 0 | 0 | 2 | 1 |
| L3a | 34 | 5 | 29 | 0 | 0 | 3 | 0 | 0 | 2 | 1 |
| L3b | 34 | 5 | 29 | 0 | 0 | 3 | 0 | 0 | 2 | 1 |
| C5a | 31 | 5 | 26 | 1 | 0 | 3 | 0 | 0 | 1 | 2 |

**Table S3.** Summary of SNPs (single nucleotide polymorphisms) in the genomes of evolved P. aeruginosa COPD isolates compared to corresponding initial isolates. Syn, synonymous. Nonsyn, no<u>nsynonymous.</u>

| Sample name | Total SNP | Syn | Nonsyn | Total CDS_SNP | Intergenic SNP | Start syn | Start nonsyn | Stop syn | Stop nonsyn | Premature stop |
| --- | --- | --- | --- | --- | --- | --- | --- | --- | --- | --- |
| W1b | 23 | 2 | 9 | 11 | 12 | 0 | 0 | 0 | 0 | 0 |
| W1c | 54,644 | 37,152 | 9,708 | 46,902 | 7,742 | 5 | 18 | 20 | 8 | 27 |
| W2a | 12 | 1 | 10 | 12 | 0 | 0 | 0 | 0 | 0 | 1 |
| W2b | 7 | 1 | 5 | 6 | 1 | 0 | 0 | 0 | 0 | 0 |
| W2c | 17 | 3 | 10 | 15 | 2 | 0 | 0 | 0 | 1 | 1 |
| W3a | 54,045 | 36,890 | 9,613 | 46,538 | 7,507 | 6 | 14 | 19 | 7 | 24 |
| W4a | 54,644 | 37,152 | 9,707 | 46,901 | 7,743 | 5 | 18 | 20 | 8 | 27 |
| W4b | 17 | 2 | 5 | 7 | 10 | 0 | 0 | 0 | 0 | 0 |
| W4c | 9 | 0 | 5 | 5 | 4 | 0 | 0 | 0 | 0 | 0 |
| W5a | 9 | 2 | 5 | 7 | 2 | 0 | 0 | 0 | 0 | 0 |
| W5b | 54,030 | 36,889 | 9,612 | 46,537 | 7,493 | 6 | 14 | 19 | 7 | 25 |
| W5c | 23 | 4 | 9 | 14 | 9 | 0 | 0 | 0 | 1 | 0 |
| W6a | 30 | 6 | 7 | 14 | 16 | 0 | 0 | 0 | 1 | 0 |
| W6b | 54,639 | 37,144 | 9,711 | 46,897 | 7,742 | 5 | 18 | 20 | 8 | 27 |
| W6c | 7 | 1 | 5 | 6 | 1 | 0 | 0 | 0 | 0 | 0 |
| W7d | 55,890 | 37,850 | 9,964 | 47,863 | 8,027 | 6 | 17 | 19 | 10 | 32 |
| W7e | 10 | 2 | 6 | 9 | 1 | 0 | 0 | 0 | 0 | 1 |
| W7f | 54,031 | 36,889 | 9,612 | 46,537 | 7,494 | 6 | 14 | 19 | 7 | 25 |
| W8a | 16 | 4 | 8 | 13 | 3 | 0 | 0 | 0 | 0 | 1 |
| M2a | 3 | 2 | 1 | 3 | 0 | 0 | 0 | 0 | 0 | 0 |
| M3a | 18 | 8 | 6 | 14 | 4 | 0 | 0 | 0 | 0 | 0 |
| M4a | 12 | 5 | 1 | 7 | 5 | 0 | 0 | 0 | 0 | 1 |
| M5a | 37 | 2 | 2 | 4 | 33 | 0 | 0 | 0 | 0 | 0 |
| L2a | 29,004 | 18,616 | 5,856 | 24,520 | 4,484 | 4 | 17 | 10 | 7 | 24 |
| L2b | 28,997 | 18,609 | 5,858 | 24,515 | 4,482 | 4 | 17 | 10 | 7 | 24 |
| L3a | 29,008 | 18,611 | 5,864 | 24,523 | 4,485 | 4 | 17 | 10 | 7 | 24 |
| L3b | 28,997 | 18,606 | 5,861 | 24,515 | 4,482 | 4 | 17 | 10 | 7 | 24 |
| C5a | 33,170 | 21,769 | 6,716 | 28,547 | 4,625 | 2 | 11 | 22 | 7 | 26 |

## Captions for Supplementary Datasets

**Supplementary Dataset S1.** KEGG terms enriched by significantly down-and up-regulated genes in evolved *P. aeruginosa* COPD isolates compared to their corresponding initial isolates (*p* < 0.05).

**Supplementary Dataset S2.** Expression of QS-activated and virulence-related genes in evolved *P. aeruginosa* COPD isolates compared to their corresponding initial isolates. Data shown are log2 fold change, *p* < 0.05.

